# Metabolic plasticity of T-cell therapies: multi-omic profiling of interacting human tumor-infiltrating lymphocytes and autologous tumor adoptive cell therapy

**DOI:** 10.1101/2022.09.28.509590

**Authors:** Melisa Martinez-Paniagua, Cara Haymaker, Jonathan Robinson, Michal Harel, Caitlin Creasy, Jay R T. Adolacion, Xingyue An, Mohsen Fathi, Ali Rezvan, Monish Kumar, Amit Amritkar, Scott E. Woodman, Rodabe N. Amaria, Tamar Geiger, Patrick Hwu, Chantale Bernatchez, Navin Varadarajan

**Author notes:** Corresponding author: Navin Varadarajan Address: William A. Brookshire Department of Chemical and Biomolecular Engineering, University of Houston, Houston, Texas 77204, USA.

## Abstract

Adoptive cell therapy (ACT) based on *ex vivo* expanded autologous tumor-infiltrating lymphocytes (TILs) can mediate durable antitumor responses even in heavily pretreated patients. However, only a subset of patients responds to ACT; efforts to identify correlates of response have focused on profiling the tumor or the TIL but rarely in an interactive environment. Interactive profiling can provide unique insights into the clinical performance of TILs since the fate, function, and metabolism of TILs are influenced by autologous tumor-derived factors. Here, we performed a suite of cell-sparing assays dubbed holistic analysis of the bioactivity of interacting T cells and autologous tumor cells (HABITAT). HABITAT profiling of TILs used for human ACT and their autologous tumor cells included function-based single-cell profiling by timelapse imaging microscopy in nanowell grids (TIMING); multi-omics using RNA-sequencing and proteomics; metabolite inference using genome-scale metabolic modeling, and pulse-chase assays based on confocal microscopy to profile the uptake and fate of fatty acids (FA). Phenotypically, the ACT TILs from both responders (Rs) and nonresponders (NRs) were comprised of predominantly effector memory T cells (T_EM_ cells) and did not express a high frequency of programmed death ligand-1 (PD-L1) and showed no differences in TCR diversity. Our results demonstrate that while tumor cells from both Rs and NRs are efficient at uptaking FAs, R TILs are significantly more efficient at utilizing FA through fatty acid oxidation (FAO) than NR TILs under nutrient starvation conditions. While it is likely that lipid and FA uptake is an inherent adaptation of TIL populations to lipid-rich environments, performing FAO sustains the survival of TILs and allows them to sustain antitumor cytolytic activity. We propose that metabolic plasticity enabling FAO is a desirable attribute of human TILs for ACT leading to clinical responses.

## INTRODUCTION

Successful cancer immunotherapy, such as the adoptive T-cell therapy (ACT), chimeric antigen receptor (CAR), or immune checkpoint blockade (ICB), relies on T-cells that can infiltrate the tumor and unleash potent antitumor functions, including direct cytolysis and cytokine secretion (Grasso *et al*., 2020; Krishna *et al*., 2020; Wei *et al*., 2019). In clinical trials, ACT has shown 30-40% durable antitumor regression in pretreated melanoma patients (Andersen *et al*., 2016; Goff *et al*., 2016; Seitter *et al*., 2021). Although striking responses have been seen in individual patients, translating the success of expanded tumor-infiltrating lymphocyte populations (TILs) to ACT for cancers other than melanoma is challenging (Stevanovic *et al*., 2015). The recognition that reactivity to neoantigens arising from widely mutated tumor suppressors like p53 exists in TIL and can be harnessed to focus the anti-tumor response has expanded the clinical promise of TIL-ACT to epithelial cancers like breast, colon, and pancreas (Malekzadeh *et al*., 2019; Tran *et al*., 2016; Zacharakis *et al*., 2022).

Expanded TILs for ACT comprise a heterogenous collection of T-cells with variation in antigen-specificity, phenotype and differentiation status, and persistence. Advances in single-cell ribonucleic acid (RNA)-sequencing (scRNA-seq), mass cytometry, and T-cell receptor (TCR) repertoire profiling have identified several T-cell biomarkers potentially correlated with clinical responses. For example, we have previously reported that the expression of BTLA (B-and-T lymphocyte-attenuator) on CD8^+^ TIL in the infusion product enables serial killing and survival after infusion and is associated with favorable clinical outcomes (Haymaker *et al*., 2015). Independently, scRNA-seq of the dominant clonotypes with a single patient-derived TIL targeting the KRAS (G12D) hotspot mutation showed *IL7R, ITGB1, KLF2*, and *ZNF683* are enriched in TILs that persist after infusion (Lu *et al*., 2019). In addition, profiling of human TIL products used to treat metastatic melanoma showed that the frequency of a progenitor T-cell population (CD39^-^CD69^-^) with self-renewal characteristics was shown to mediate long-term tumor control and T-cell persistence (Krishna *et al*., 2020). These studies are consistent with the preclinical paradigm that younger, less differentiated cells are capable of more robust antitumor responses but paint an incomplete picture since highly differentiated TILs can induce durable tumor regressions following ACT (Gattinoni *et al*., 2011; Maus *et al*., 2014).

The success of TIL-ACT/immunotherapy is also shaped by a variety of tumor-intrinsic factors, including the expression of programmed death ligand-1 (PD-L1) protein, interferon-responsive signatures, tumor mutational burden, and presentation of neoantigens (Ayers *et al*., 2017; Lauss *et al*., 2017; Tokito *et al*., 2016). More recently, there has been an increased appreciation for the role of metabolism within tumor cells and the tumor microenvironment (TME) and how this can impact T-cell function and fate. For example, proteomic profiling demonstrated that high oxidative-phosphorylation and lipid metabolism in tumor cells was associated with the response in advanced-stage melanoma patients treated with ACT and ICB (Harel *et al*., 2019). Preclinically, it has been shown that tumor/TME-derived lactate, cholesterol, and free fatty acids negatively impact T-cell function and fate (Cascone et al., 2018; Ma et al., 2019b; Xu et al., 2021b).

Although these studies have helped advance our understanding of responders of TIL-ACT, there are no studies that profile human TILs in the context of autologous pretreatment tumors in an interactive environment. A contextual and interactive study is an essential feature for understanding the performance of TILs since, as highlighted above, the fate, function, and metabolism of TILs are influenced by matched tumor-derived factors. We, therefore, profiled a set of 16 patients for whom we had access to the TILs and the autologous/matched tumors. Our results from a suite of holistic analyses of the bioactivity of interacting T-cells and autologous tumor cells (HABITAT) assays: dynamic single-cell tumor-cell functional assays; RNA-seq and proteomics upon co-culture; and genome-scale modeling illustrate the metabolic plasticity of TILs to perform fatty acid oxidation (FAO) under nutrient limitation sustains killing function leading to potent anti-tumor efficacy.

## RESULTS

### Patient characteristics and overall response to TIL therapy

We used a set of sixteen samples from metastatic melanoma patients treated under TIL infusion protocols, with eight responders (Rs) and eight nonresponders (NRs). We chose these sixteen samples based on the availability of autologous tumor lines, overall responses, and a high frequency of CD8^+^ T-cells. **Table 2** describes the patient characteristics, mutational status of the tumor, and an overview of the cellular infusion products for all 16 patients (**Figure S1**). The median duration of response was 19.3 months for NR (range 3-22 months) and 39 months for the R (range 9 months-6 years), with responses that were stable for at least four years in 4/8 of these patients (**Figure S2**).

**Table 1.** List of abbreviations used in this study

|  |  |
| --- | --- |
| 2NBDG | Two-(N-(seven-Nitrobenz-two-oxa-one, three-diazol-four-yl)amino)-two-deoxyglucose |
| ACC1 | Acetyl-CoA carboxylase one |
| ACT | Adoptive cell therapy |
| AICD | Activation-induced cell death |
| ATCC | American type culture collection |
| BTLA | B-and-T lymphocyte-attenuator |
| CAR | Chimeric antigen receptor |
| CM | Control media |
| CoA | Coenzyme A |
| CPT1a | Carnitine palmitoyltransferase one alpha |
| CTC | Circulating tumor cell |
| DE | Differential expression |
| DEG | Differentially expressed gene |
| DEP | Differentially expressed protein |
| EMT | Epithelial to mesenchymal transition |
| FA | Fatty acid |
| FAO | Fatty acid oxidation |
| FASN | Fatty acid synthase |
| FDA | Food and drug administration |
| FDR | False discovery rate |
| FIBP | Fibroblast growth factor intracellular-binding protein |
| GEM | Genome-scale metabolic model |
| GPX4 | Glutathione peroxidase four |
| GSEA | Gene set enrichment analysis |
| HABITAT | Holistic analysis of the bioactivity of interacting T cells and autologous tumor cells |
| HBSS | Hank's balanced salt solution |
| HLA | Human leukocyte antigens |
| HMR | Human metabolic reaction |
| ICB | Immune checkpoint blockade |
| IFNγ | Interferon gamma |
| IL | Interleukin |
| IND | Investigational new drug |
| IRB | Institutional review board |
| LD | Lipid droplet |
| MART-1 | Melanoma antigen recognized by T cells one |
| MSigDB | Molecular signatures database |
| MTG | MitoTracker green |
| MTR | MitoTracker red |
| NR | Nonresponder |
| OA | Oleic acid |
| PBL | Peripheral blood lymphocyte |
| PD1 | Programmed cell-death protein 1 |
| PDA | Pancreatic ductal adenocarcinomas |
| PDL1 | Programmed death ligand 1 protein |
| PGC1α | Peroxisome proliferator-activated receptor-gamma coactivator one alpha |
| PPAR | Peroxisome proliferator-activated receptor |
| REP | Rapid expansion protocol |
| RNA-seq | RNA sequencing |
| R | Responder |
| scRNA-seq | Single-cell RNA-seq |
| SIRM | Stable isotope-resolved metabolomics |
| tContact | Time of cumulative conjugation after the establishment of a synapse |
| TCR | T cell receptor |
| tDeath | Time to induce tumor cell apoptosis |
| T<sub>EM</sub> | Memory T cells |
| T<sub>ex</sub> | Exhausted T cells |
| TGFβ | Transforming growth factor beta |
| TILs | Tumor-infiltrating lymphocyte cells |
| TIMING | Timelapse imaging microscopy in nanowell grids |
| tINIT | Task-driven integrative network inference for tissues |
| TME | Tumor microenvironment |
| T<sub>PEX</sub> | Progenitor exhausted like T cell |
| T<sub>reg</sub> | Regulatory T cells |
| Tres | Tumor-resilient T cells |
| T<sub>RM</sub> | Tissue-resident memory T cells |
| tSeek | Time to establish a synapse |

**Table 2.** Clinical and demographic characteristics of the patients and their TILs/tumors profiled in this study.

| TIL Number | Number of TIL infused (x10<sup>9</sup>) | Progression on anti-PD-1 therapy before TIL therapy (Yes/No) | Progression on Ipi before TIL therapy (Yes/No) | Progression on Ipi Before TIL Harvest (Yes/No) | Progression on HD IL-2 before TIL | Progression on BRAF inhibitor before TIL (does not include sorafenib) | Progression on Biochemo before TIL (does not include chemo alone) | Number of systemic therapies before TIL | Gender | Best iRC | Response | Age | BRAF mutation | NRAS mutation | Kit mutation |
| --- | --- | --- | --- | --- | --- | --- | --- | --- | --- | --- | --- | --- | --- | --- | --- |
| 3140 | 1.54 | Yes | Yes | Yes | No | No | No | 8 | Female | SD | NR | 60 | mutated | WT | WT |
| 2800 | 292 | No | No | Yes | No | Yes | No | 10 | Male | PD | NR | 31 | V600E | WT | WT |
| 2765 | 143.8 | Yes | Yes | Yes | Yes | No | No | 6 | Male | SD | NR | 34 | WT | WT | WT |
| 2559 | 69.5 | No | No | N/A | Yes | Yes | No | 12 | Male | SD | NR | 34 | V600E | NT | NT |
| 2458 | 14.30 | No | No | N/A | No | No | Yes | 1 | Male | PD | NR | 45 | V600E | WT | WT |
| 2453 | 69 |  |  |  | Yes |  |  | 1 | Male | SD | NR | 63 | V600E | Q61K | M541L |
| 2399 | 89 | No | No | No | Yes | No | Yes | 6 | Male | SD | NR | 52 | WT | Q61R | WT |
| 2373 | 59.20 | No | No | N/A | Yes | No | No | 2 | Male | SD | NR | 50 | V600E | WT | WT |
| 2576 | 106.00 | Yes | No | No | Yes | No | Yes | 6 | Male | PR | R | 65 |  |  |  |
| 2508 | 27.30 | No | Yes | Yes | No | No | No | 2 | Female | CR | R | 50 | V600E | WT | WT |
| 2482 | 55.00 | No | No | N/A | Yes | No | No | 4 | Female | PR | R | 41 | V600E | WT | WT |
| 2427 | 16.3 |  |  |  | N/A |  |  | 0 | Female | PR | R | 54 | V600E | NT | NT |
| 2379 | 100.00 | No | No | N/A | Yes | No | No | 1 | Female | CR | R | 46 | V600E | WT | NT |
| 2364 | 107 |  |  |  | Yes |  |  | 1 | Male | CR | R | 18 | No | Q61K | No |
| 2357 | 109.00 | No | No | N/A | Yes | Yes | No | 5 | Male | CR | R | 53 | K601E | WT | WT |
| 2340 | 46 |  |  |  | No |  |  | 1 | Female | PR | R | 48 | V600E | WT | WT |

### R-TILs had improved survival compared to NR-TILs

Although we had reported in prior studies an association between the number of TIL infused and the percentage of CD8^+^ TIL in the infusion product with the outcome of TIL therapy, this difference was not significant in an updated cohort of patients who progressed on checkpoint blockade therapy before being treated with TIL therapy (**Table 2**) (Forget *et al*., 2018). As we have published previously, most TILs utilized for this study were of the T_EM_ phenotype (CCR7^+^CD45RA^-^) (Forget *et al*., 2018). In addition, within the CD8^+^ TILs, we evaluated CD28 expression (less differentiated cells) or PD1 expression (exhausted cells) and observed no significant differences between R and NR samples (**Figure S3**) (Prieto *et al*., 2010; Thommen *et al*., 2018).

We next investigated if the persistence of specific subsets of TILs after infusion can indicate a response to therapy. We utilized blood samples longitudinally collected following TIL infusion to track the T-cell clonotypes in the peripheral blood. To identify TCR sequences derived from the TIL products potentially associated with T-cells recognizing the tumor, expanded TILs were co-incubated with autologous tumor cells and flow-sorted into CD8^+^CD69^+^ (activated, tumor-specific) and CD8^+^CD69^-^ (unactivated) T-cell subsets. We determined the TCRβ CDR3 sequences belonging to the two separate subsets by high-throughput sequencing to track T-cell clonotypes (Spindler *et al*., 2020). There was no consistent pattern between the number of CD69^+^ or CD69^-^ CD8^+^ T-cell clones in the infusion product or their respective persistence in the blood over time that delineated clinical outcome (**Figure S4**).

Since we established that the phenotype and clonal selection were not showing a clear pattern between the CD69 expression and persistence between R and NR TILs, we next investigated the functional capacity of the TILs. We performed paired assays to map the functional capacity wherein each TIL population was evaluated for function against autologous tumor cells. We confirmed that the TILs were reactive to the autologous tumors by interferon γ (IFNγ) ELISA upon co-culture in a human leukocyte antigen (HLA)-restricted manner (**Figure S5**). As our first HABITAT assay, we mapped the kinetics of the interaction of TIL with autologous tumor cells in a tumor-killing event using a high-throughput single-cell technology <u>T</u>imelapse Imaging <u>M</u>icroscopy In <u>N</u>anowell <u>G</u>rids (TIMING) [**Figure 1A-C**]. Kinetic measurements, including the time to establish a synapse (tSeek), time of cumulative conjugation after the establishment of a synapse (tContact), and the time to induce tumor cell apoptosis (tDeath) were not different for TILs derived from either R or NR (**Figure S6A-C**). The number of cells that established a synapse was also not different between R and NR TILs (**Figure S6D**). We quantified the persistent motility (directional motility for at least one cell diameter) for the TILs and tumor cells within the TIMING assays. We observed that R TILs showed more motility and polarization than NR TILs, enabling more efficient synapse formation (**Figure 1C-D**). By contrast, the NR tumor cells, as opposed to R tumor cells, showed increased motility and polarization with and without conjugation to autologous TILs, (**Figure 1E**). Lastly, we compared the frequency of TILs undergoing activation-induced cell death (AICD) and observed that despite a comparable frequency of synapse formation, NR TILs underwent AICD at a higher frequency than R TILs (**Figure 1F**). Collectively, the results from the TIMING assays established that R TILs are functionally equivalent but less prone to AICD than NR TILs. Regarding the tumor cells, NR tumor cells were more motile than R tumor cells, which could facilitate escape from T-cell recognition.

**Figure 1.**
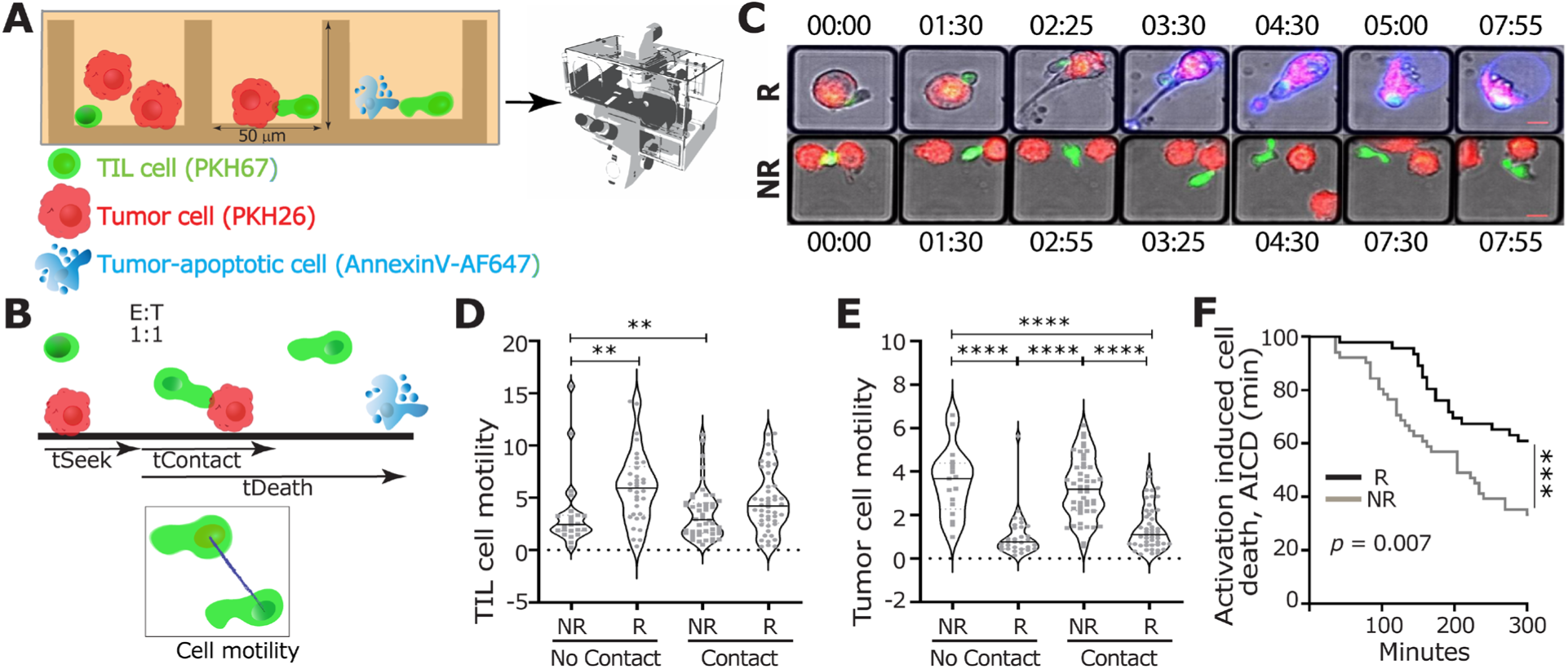
TILs from responders showed improved motility and survival. **(A)** Timelapse Imaging Microscopy In Nanowell Grids (TIMING) co-incubates thousands of effectors (TILs) and target cells (autologous tumor cell lines) in nanowell arrays. The entire nanowell array is imaged using automated epifluorescent microscopy to track the interactions between TIL and tumor cells with single-cell resolution. **(B)** Quantitative metrics used to describe the interactions of the TILs with their autologous tumor cells. The cell motility is measured using the average displacement of the TIL or tumor centroid during three consecutive time points (bottom). **(C)** Representative composite micrographs illustrating the interaction between R or NR TILs and corresponding autologous tumor cells within nanowells. Images were representative of four different experiments. Scale bar = 10 μm, the time displayed as “hh:mm” where “h” represents hours and “m” minutes. Arrays were imaged for 8 hours at an interval of six minutes. (**D-E**) Comparison of the motility of individual TILs (D) or autologous tumor cells (E) with (contact) and without conjugation (no contact). (**F**) Survival curve of TILs that induced death of target cells. All data in panels D-F are from evaluating a total of 1,818 1E:1T events with evidence of synapse formation (conjugation) within nanowells. *p* values for comparisons were computed using a non-parametric Mann-Whitney test. (n = 4, two Rs and two NRs). \**p* < 0.05, \*\**p* < 0.01,\*\*\*\**p* < 0.0001 in D-F.

### NR tumor cells have features of mesenchymal/amoeboid motility

To identify the mechanistic basis for the increased motility of NR tumor cells, we performed RNA-seq on the tumor cells (**Figure 2A**). The tumor mutational status evaluation indicated that the tumors in both groups were wild-type *NRAS* and *c-Kit* but mutated *BRAF* (**Table 2**). To identify differentially expressed pathways, we performed gene set enrichment analysis (GSEA), clustered them based on the overlap of genes, and visualized them using Cytoscape (**Figure 2B**). We identified five significant clusters showing pathways enriched in NR tumor cells: one cluster of pathways related to cellular metabolism, another cluster related to cancer cell metastasis, and three clusters related to cellular motility, adhesion, and migration (**Figure 2B**). The cluster of pathways associated with metabolism was primarily enriched in pathways associated with the utilization of fatty acids and adipogenesis (**Figure 2B**). The pathways related to metastasis were predominantly comprised of pathways related to epithelial-to-mesenchymal transition (EMT). Since melanoma is a non-epithelial tumor, it does not undergo canonical EMT, and the acquisition of EMT-like signatures has been associated with increased stem-cell and neural-crest (highly migratory) properties, and amoeboid migration (**Figure 2C**) (Rodriguez-Hernandez *et al*., 2020; Sanz-Moreno *et al*., 2011). The three clusters of migration-related pathways were enriched in negative regulation of epithelial cell proliferation and migration and increased amoeboid migration (**Figure 2C**). Based on a published signature of ameboid and epithelial morphology of A375 melanoma cells, we also confirmed that NR tumor cells in our cohort are ameboid-like (**Figure S7**)(Georgouli *et al*., 2019). One of the mechanisms of immunosuppression adopted by amoeboid A375M2 cells is via the secretion of soluble factors like transforming growth factor (TGFβ) and interleukin 10 (IL-10)(Georgouli *et al*., 2019). GSEA comparisons confirmed that NR tumor cells enriched the same immunomodulatory factors that induce tumor-promoting macrophages (**Figure 2D**).

**Figure 2.**
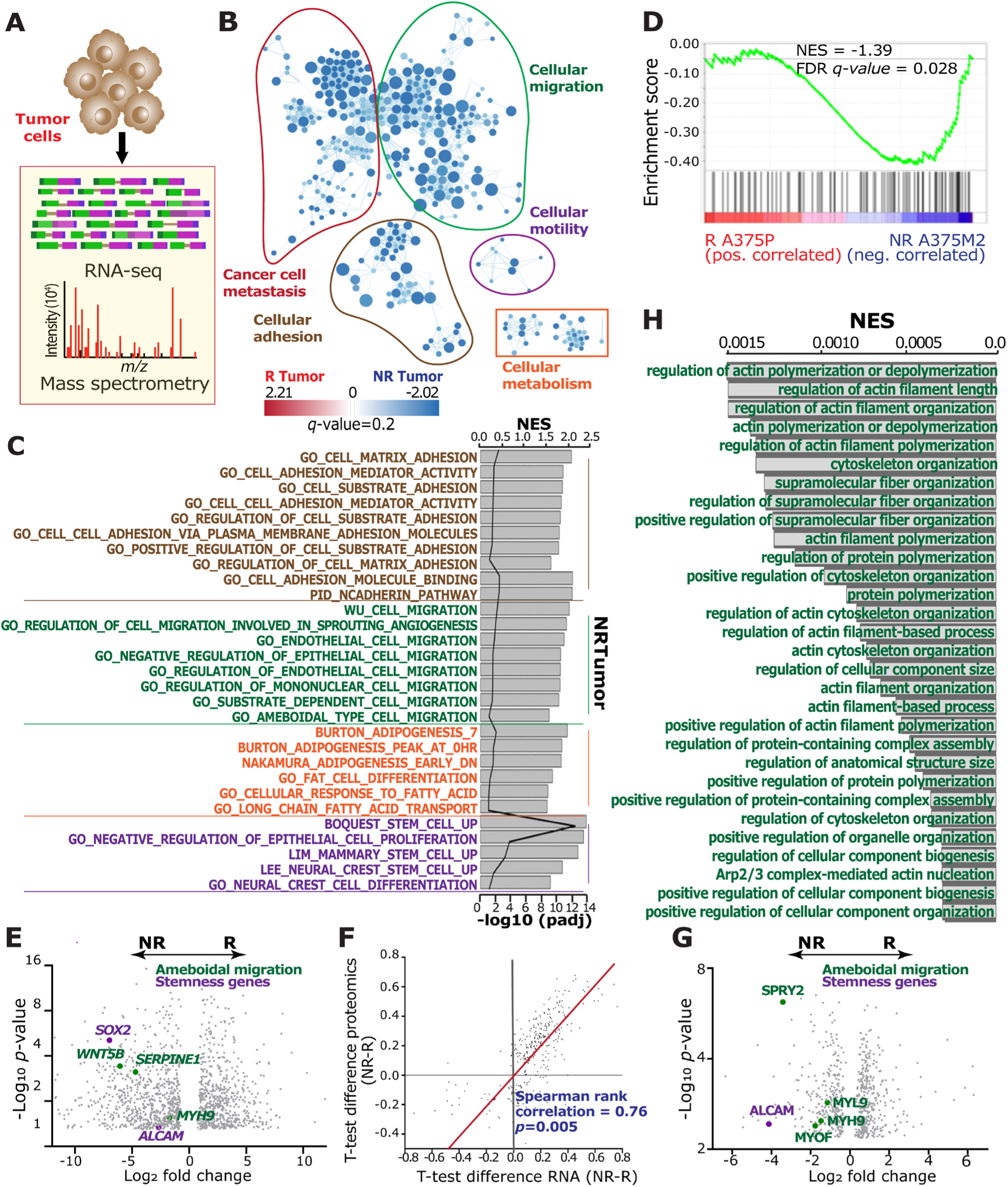
NR tumor cells are migratory with amoeboid-like characteristics. **(A)** Overview of tumor cell profiling by RNA-seq and mass spectrometry. **(B)** The gene set enrichment analysis derived enriched C2 curated pathways in three R tumor and three NR tumor cells plotted in Cytoscape using a cutoff false discovery rate (FDR) *q* value of 0.01 and *P-*value of 0.005. Clusters of pathways are labeled as groups with a similar theme as indicated. All pathways enriched in NR tumor cell populations. **(C)** List of pathways enriched on the NR tumor cells grouped by color. Brown: cell adhesion pathways; green: amoeboid migration pathways; orange: metabolism-related pathways and purple: stemness genes. **(D)** Gene-set enrichment analysis plots positively correlate with amoeboid signature with NR tumor cells. **(E)** Volcano plot generated using the mRNA data fold change (x-axis) and *p-*values of DEGs (y-axis) showing differences between R and NR tumor cells of the genes. Genes of interest related to the pathways described in C are annotated. **(F)** 2D enrichment plot showing a positive correlation of mRNA data (x-axis) and proteomics data (y-axes) results (Spearman rank correlation = 0.76, *p* = 0.005). **(G)** Volcano plot of the DEPs as a function of the log2 difference between the averages of the two response groups (x-axis) and the -log10 *p*-value (y-axis). DEPs are colored in gray. Proteins related to ameboid migration and stemness genes are highlighted. **(H)** Pathways related to ameboid-like migration of tumor cells enriched in NR tumor cells (n = 6, three Rs, and three NRs tumor cells).

Myosin II primarily controls amoeboid migration of cancer cells at the molecular level, leading to hyper-activation of Rho-ROCK1/2 signaling and non-canonical Wnt signaling (Rodriguez-Hernandez *et al*., 2020; Sahai and Marshall, 2003). Therefore, the candidate differentially-expressed genes (DEGs) comparing the NR tumor cells to the R tumor cells were consistent with the pathway analyses: genes associated with amoeboid migration (*MYH9*, heavy chain of Myosin II; *WNT5B* and *SERPINE1*), and stemness genes (*ALCAM* and *SOX2*) were enriched in NR tumor cells (**Figure 2E, Table 6**).

We performed unbiased proteomics on these tumor cells to validate these observations. Overall, there were no major differences in detected proteins between the R and NR tumor cells (**Figure S8**). After filtering, normalization, and imputation, we identified 7087 proteins in at least 50% of the samples. The NR tumor cells clustered together, and these were separated from the group of R tumors. We identified 250 differentially expressed proteins (DEPs) with higher expression in R tumor cells and 218 DEPs with higher expression in NR tumor cells (FDR<0.05; **Figure S8**). To identify pathways significantly enriched in both the RNA-seq and proteomics datasets, we used 2D enrichment, as we have previously published (Harel *et al*., 2019). There was a strong correlation between the RNA-seq and proteomic pathways (**Figure 2F**, Spearman correlation 0.76). At the molecular level, the DEPs based on proteomics data validated proteins related to amoeboid motility (MYH9, MYL9, MYOF, and SPRY2) and stemness (ALCAM) (**Figure 2G, Table 7**). In addition, the migration-related pathways were enriched in positive regulation of actin filament and cytoskeleton organization (**Figure 2H**).

In aggregate, NR tumor cells have mesenchymal/amoeboid motility features when evaluated using functional and morphological assays (TIMING), transcriptional signatures (RNA-seq), and proteomic profiling similar to those reported previously (Rodriguez-Hernandez *et al*., 2020; Sanz-Moreno *et al*., 2011).

### RNA-seq identifies that R TILs upregulate fatty acid oxidation when co-cultured with autologous tumor cells

We next wanted to investigate why the R TIL, compared to NR TILs, were more resistant to apoptosis when interacting with autologous tumor cells. To identify mechanisms responsible for this difference, we performed a HABITAT assay wherein we co-incubated TIL with autologous tumor cells for 12 h to promote functional interactions leading to T-cell activation (**Figure 3A**). We subsequently flow-sorted the CD8^+^ TILs into two distinct populations marked by CD69 expression and prepared cDNA libraries based on the sorted TILs (**Figures S9-S10**).

**Figure 3.**
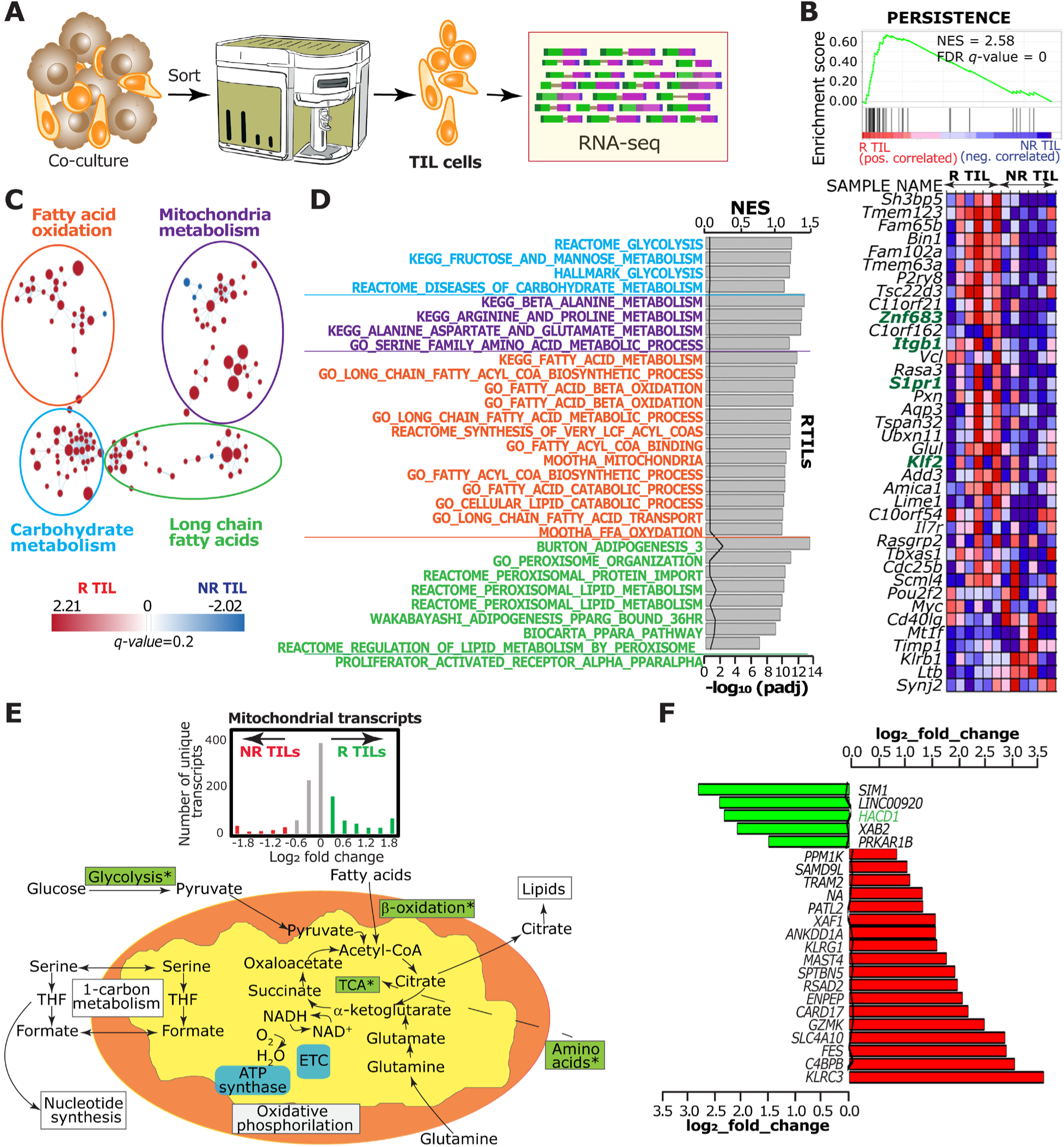
Responder TILs upregulate pathways related to T-cell persistence and mitochondrial fatty acid oxidation compared with the NR TILs. **(A)** Overall experiment design for profiling the pathways upregulated in TILs upon co-culture with autologous tumor cells (12 h). Six R TIL populations and six NR TIL populations were profiled by RNA-seq. **(B)** GSEA-based comparison of R and NR TILs to a known signature of T-cell persistence (GSE136394) (Lu *et al*., 2019). **(C)** The gene set enrichment analysis derived C2 curated pathways visualized in Cytoscape using a cutoff false discovery rate (FDR) *q* value of 0.01 and *p-*value of 0.005. Four major clusters are highlighted. All pathways are enriched in R TIL populations. **(D)** Pathways enriched in the R TILs, grouped by color. Blue: carbohydrate metabolism, purple: mitochondria metabolism, orange: fatty acid oxidation, green: long-chain fatty acid-related pathways. **(E)** Comparative assessments of the mitochondrial enrichment transcripts (MitoCarta) within R and NR TILs. Schematic depicting the mitochondrial pathways enriched (green boxes) in R TILs. **(F)** Heatmap illustrating the persistence-related genes (highlighted in green) comparing the R and NR TILs.

Since T-cell exhaustion status is known to impede TIL efficacy, we compared the signatures of R and NR TILs against published signatures of exhaustion. When tested against three different signatures of CD8^+^ T exhausted (T_ex_) and T progenitor exhausted-like (T_PEX_) cells, there was no significant difference between the R and NR TILs (**Figure S11A**) (Galletti *et al*., 2020; Khan *et al*., 2019; Thommen *et al*., 2018). Although T-cell persistence correlates to a clinical response with TIL-based therapies, T-cell persistence is influenced by antigen avidity, tumor microenvironment, and host cytokine support. We compared the R and NR TILs against a published signature of T-cell persistence based on clonotypes targeting KRAS (G12D) and confirmed that the R TILs were enriched for persistence (Lu *et al*., 2019). At the molecular level, we observed an enrichment of crucial transcription factors *ZNF683, S1pr1*, and *KLF2* within our R TILs (**Figure 3B**).

To derive a complete understanding of the differences between the R and NR TILs, we performed GSEA to identify differentially expressed pathways (false discovery rate [FDR] q-value < 0.2), clustered them based on the overlap of genes and visualized them using Cytoscape (**Figure 3C**). Four major clusters emerged showing pathways related to FAO, long-chain fatty acids, mitochondria, and carbohydrate metabolism (**Figure 3C**). The pathways in the carbohydrate metabolism cluster make up the pathways associated with glycolysis (**Figure 3D**). The mitochondrial cluster contains mainly amino acid metabolism pathways, including glutamine/glutamate metabolism, that are known to upregulate upon T-cell activation (**Figure 3D**). To directly probe enhanced mitochondrial involvement, we mapped the mitochondrial transcripts onto MitoCarta and performed GSEA. GSEA confirmed the enrichment of mitochondrial transcripts in R TIL compared to NR TILs (**Figure 3E**). The two clusters associated with lipids were directly enriched in fatty acyl-CoA biosynthesis, peroxisomal and mitochondrial beta-oxidation (FAO) [**Figure 3D**].

At the molecular level, as expected by the TILs from both groups being predominantly T_EM_, there were only 36 candidate DEGs (FDR q < 0.1) between the NR and R TILs (**Figure 3F**). Furthermore, the gene, *HACD1*, that encodes for the very-long-chain (3R)-3-hydroxyacyl-CoA dehydratase 1, was upregulated in the R TILs, consistent with the pathway-level enrichment of fatty acid utilization described above (**Figure 3F**). Finally, we confirmed that R TILs showed enrichment in signatures associated with FAO via comparison to published signatures of FAO [**Figure S11B**] (Shen *et al*., 2020). Altogether, these results suggest that R TILs upregulate pathways associated with mitochondrial oxidation and glycolytic metabolism when co-cultured with autologous tumor cells.

### Genome-scale metabolic modeling shows enrichment of fatty acid metabolites in R TILs when co-cultured with autologous tumor cells

Although stable isotope-resolved metabolomics (SIRM) is the preferred approach to analyzing the metabolic reaction pathways and networks, the limited number of TILs available from patients precluded us from performing these assays (Jang et al., 2018; Ma et al., 2019a). To overcome this limitation, we took advantage of genome-scale metabolic models (GEM) to derive a mechanistic understanding of intracellular metabolism (**Figure 4A**). To explore the metabolic component of the gene expression changes within the TILs, we used the differential expression analysis results from RNA-seq and integrated it with a human GEM (Human1) (Robinson *et al*., 2020) using the reporter metabolites method. Since we have matched RNA-seq data on TIL and the tumor cells, we extracted the relevant subset of the generic human GEM for each cell type using task-driven integrative network inference (tINIT) (Agren *et al*., 2014).

**Figure 4.**
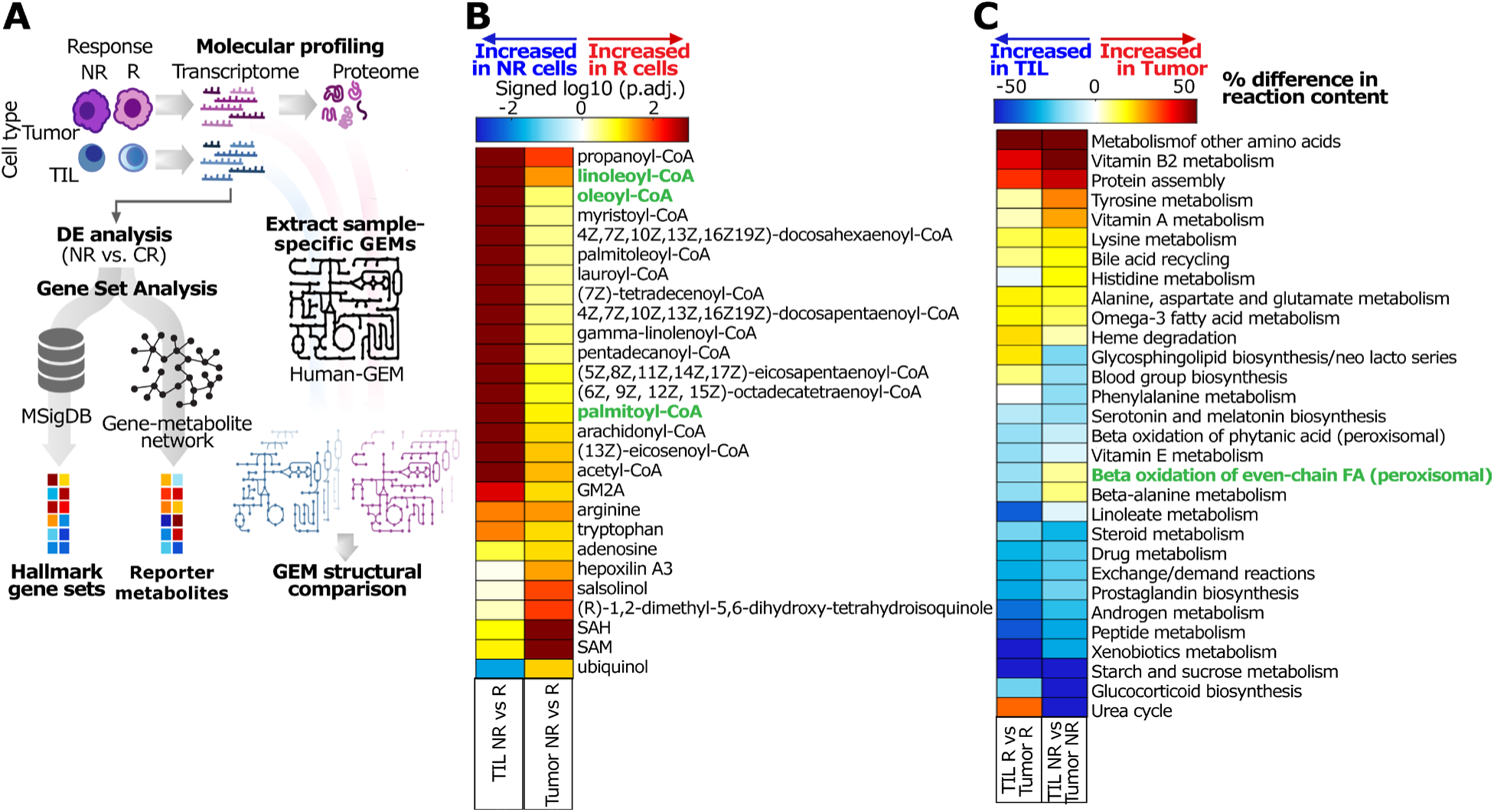
Genome-scale integrated metabolic modeling of the reaction networks in TIL and autologous tumor cells. **(A)** Model generation and data analysis workflow. Differential expression (DE) analysis was performed on the transcriptomic data within each cell type to evaluate the difference between NR and R samples. Gene set analysis was performed using the Molecular Signatures Database (MSigDB) Hallmark gene set collection or the gene-metabolite interaction network extracted from the Human1 model. Transcriptomic and proteomic data were also integrated with Human1 to extract a GEM specific to each sample, the structures of which were compared to predict differences in their active metabolic networks. **(B)** Reporter metabolite analysis results show the enrichment of gene expression changes associated with different metabolites. Only the top 15 most significant metabolites for each comparison (NR vs. R TILs and NR vs. R Tumor cells) are shown. The *p*-values are adjusted for the false discovery rate, log-transformed, and signed based on the direction of expression change: positive (red) corresponds to higher expression in R cells, whereas negative (blue) corresponds to higher expression in NR cells. **(C)** Cell-type-specific models were grouped by cell type (TIL or tumor) and response status (NR or R), and the average number of reactions in each subsystem was determined. The percent difference in the number of reactions was calculated for a difference in cell type (TIL vs. Tumor) for the same response status. For each comparison, only the top 30 subsystems exhibiting the most significant difference in reaction content are shown.

The normalized RNA-seq data from each sample was separated into four groups according to cell type (tumor or TIL) and response status (NR or R). At the level of individual metabolites, the significantly enriched metabolites within R TILs were dominated by peroxisome and mitochondria-derived fatty acyl-CoA: *e.g.,* palmitoyl-CoA, linoleoyl-CoA, and oleoyl-CoA (**Figure 4B**). Since the RNA-seq of the TIL was obtained after co-culture with the autologous tumor cells, we also investigated reaction networks enriched in the TILs compared to their autologous tumor cells. An entire set of reaction networks associated with mitochondrial beta-oxidation was enriched in the model network comparison of R TIL vs. R tumor cells but not in the corresponding NR TIL vs. NR tumor cells (**Figure 4C**). Collectively, the results from metabolic modeling studies illustrate that the ability to perform mitochondrial beta-oxidation in the presence of autologous tumor cells is a defining and differential feature of R TILs.

### R TILs uptake FA and perform mitochondrial FAO under conditions of nutrient starvation

In mouse models of melanoma and colon cancer, it has been demonstrated that T-cells undergo metabolic reprogramming within the TME, and the uptake of lipids by TILs leads to T-cell functional impairment and exhaustion (Xu *et al*., 2021b). Although our combined results from RNA-seq and genome-scale modeling support lipid utilization and mitochondrial FAO within R TILs, these T-cells were not functionally exhausted (**Figure 1**), did not show elevated expression of PD1 (**Figure S3**), or did not show transcriptional signatures of exhaustion (**Figure S11A**). We factored in two considerations to validate the prediction of R TILs performing FAO when co-cultured with tumor cells. First, TILs are a collection of heterogeneous populations (as validated with TCR Vβ clonotypes in our samples, **Figure S4**) with divergent reactivities; hence, single-cell profiling would be essential to mapping the differences in metabolism correlated with clinical response. Second, the validation needs to be performed with sample-sparing assays as the number of patient TILs available was limited. Accordingly, we performed HABITAT assays by co-culturing the TIL with the autologous tumor cells. We analyzed them using multiplexed flow cytometry and quantitative 3D confocal microscopy that offer a single-cell resolution to directly quantify glucose, lipid, and mitochondrial metabolism using metabolic dyes.

Glucose deprivation due to the microenvironment conditions and loss of mitochondrial function can impair the antitumor activity of T-cells. Thus, we investigated whether TILs before and after the co-culture with their autologous tumors altered mitochondrial metabolism (**Figure S12**). As a result, both R and NR TILs had similar mitochondria mass (assessed by MitoTracker green [MTG]), glucose uptake (measured by the analog two-(N-(seven-Nitrobenz-two-oxa-one, three-diazol-four-yl)amino)-two-deoxyglucose [2-NBDG]), and mitochondrial membrane potential (measured by MitoTracker red CMX Ros [MTR]) after co-culture with the autologous tumor cells (**Figure S13**).

In order to survive a nutrient-restricted microenvironment, TIL and tumor cells must reprogram their metabolism and catabolize lipids to sustain their functions (Carracedo *et al*., 2013; Ma *et al*., 2018; Patsoukis *et al*., 2015; Pearce *et al*., 2009). To determine if the TILs uptake lipids, we measured lipids contained in the lipid droplets (LD) using Bodipy 493/503, a lipophilic fluorescent probe that recognizes polar and neutral lipids. When the TILs were co-cultured with autologous tumor cells for 12 hours, we found no differences in the LD content between R and NR TILs under standard nutrient conditions (**Figure 5A, 5B**). To map the impact of nutrient starvation on LD turnover, we set up a pulse-chase assay. We stimulated the TILs with oleic acid (OA) to increase the lipids uptake and storage and subsequently co-cultured the TILs with their respective autologous tumor cells in Hank’s balanced salt solution (HBSS) [nutrient deficient, chase] for six hours, followed by LD tracking by confocal microscopy (**Figure 5C**). There was no difference in the number of LDs between R and NR TILs upon stimulation with OA, suggesting that FA uptake was conserved in TILs (**Figure 5E**). However, upon co-incubation with autologous tumor cells under nutrient limitation conditions, the LDs pool was significantly degraded in R TILs (**Figures 5D and 5E**). These single-cell data suggest that R TILs catabolize lipids under starvation conditions while NR TILs do not.

**Figure 5.**
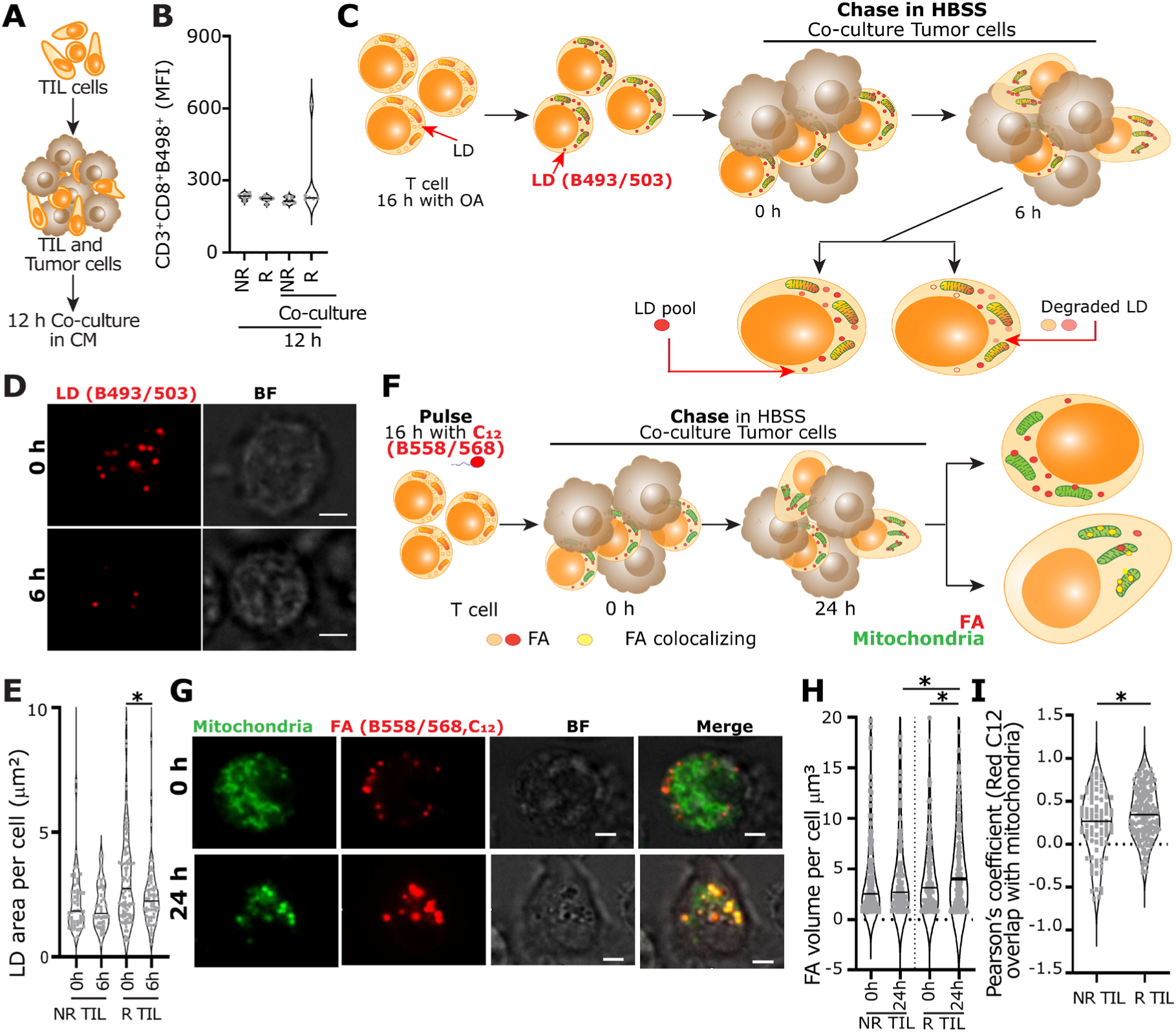
R TILs accumulate FA and perform FAO. **(A)** Monitoring the LDs in TILs. R and NR TILs were co-cultured for 12 h with their respective autologous tumor cells in complete media (CM). After co-culture, TILs were stained for LD using BODIPY 493/503 (B493/503) and analyzed by flow cytometry. **(B)** Comparison of the CD3^+^CD8^+^B493/503^+^ cells between R and NR TILs (n=12, six Rs, and six NRs). **(C)** LD pulse-chase assay. TILs were cultured in the presence of oleic acid (OA) for 16 hours to stimulate the uptake of FAs. The TILs were subsequently stained for mitochondria (MitoTracker Far Red) and B493/503 and co-incubated with autologous tumor cells under starvation conditions (HBSS). The cells were imaged using a Nikon spinning disk confocal microscope. **(D)** Representative confocal images of LDs on R TILs at zero and six hours. Scale bar = 10 μm. BF stands for brightfield, and LD for lipid droplets. **(E)** Comparative assessment of the LD area between the R TIL and NR TILs before and after co-culture with autologous tumor cells (n = 7, three R, and four NR TILs). These data are from a total of 248 individual TILs. **(F)** FA pulse-chase assay. TILs were pulsed with Bodipy 558/568 (C_12_) for 16 hours. Then, the TILs were stained using MitoTracker green to visualize the mitochondria and plated on a 96-well plate on starving conditions (HBSS) or complete media (CM) in co-culture with their respective autologous tumor cells. Finally, TILs were chased under a confocal spinning disk microscope at the beginning of the co-culture (zero) and after 24 hours. **(G)** Representative confocal images of fatty acid and mitochondria on an R TIL at the beginning of the co-culture (zero) and after 24 hours. Scale bar = 10 μm. **(H)** Comparison of the FA volume between R and NR TILs before and after co-culture with autologous tumor cells. These data are from a total of 632 individual TILs. **(I)** Comparison of the Pearson’s coefficient for the FA overlap and mitochondria between R and NR TILs after co-culture with autologous tumor cells. These data are from a total of 272 individual TILs. *p* values for comparisons were computed using a non-parametric Mann-Whitney test (n = 7, three R, and four NR cells). \**p* < 0.05, \*\**p* < 0.01,\*\*\*\**p* < 0.0001 in E,H-I.

To directly track the uptake and fate of FAs at the molecular level, we set up a second pulse-chase assay (**Figure 5F**). We pulsed the TILs with BODIPY 558/568 (C_12,_ a FA analog) [molecular length equivalent to an 18-carbon FA] for 12 hours to enable uptake of this FA (pulse). Then, we co-cultured the TILs with their autologous tumors in HBSS (starvation, chase) and tracked the total FA and their localization by confocal microscopy (**Figure 5F**). Compared to NR TILs, we found that R TILs had an increased volume of FA per cell (**Figure 5H**) and a significant colocalization of FA and mitochondria (indicative of beta-oxidation in the mitochondria, **Figure 5G-I and S14**) after 24 hours. To investigate the role of the tumor cells in influencing the uptake and utilization of FAs by the TILs, we repeated the pulse-chase assays with BODIPY 558/568 (C_12_) with just the tumor cells (no TILs added) (**Figure S15A**). Both R and NR tumor cells efficiently take up the FA (**Figure S15B**), and not surprisingly, the FA volumes per cell are much higher than TILs. Upon starvation, R and NR tumor cells utilize a fraction of these FAs via oxidation in the mitochondria (**Figure S15C**), with no significant differences between the two groups. The smaller Pearson’s coefficient for the overlap of the mitochondria and FA in tumor cells, compared to TILs, is likely because tumor cells are very efficient at FA uptake and storage, and not all of the FA is utilized immediately (**Figure S15D**). Collectively, these results directly demonstrate that while the tumor cells from both R and NR can efficiently uptake FAs and perform FAO, R TILs, when co-incubated with autologous tumors, perform FAO more efficiently than NR TILs under identical conditions.

We utilized TILs targeting a single defined epitope to directly link FAO in supporting TIL function/survival in a defined context. Accordingly, we utilized two clonal TIL lines derived from NRs that recognize the melanoma antigen recognized by T-cells 1 (MART-1), which are HLA-A*0201 restricted. We set up a kinetic cytotoxicity assay with MART-1 specific TILs as effectors, and Mel526 (MART-1 positive) tumor cells as targets in HBSS supplemented with BSA-palmitate as the source of FAs (**Figure 6A**). To modulate FAO, we used small molecules that either inhibit (etomoxir) or activate (baicalin) the rate-limiting enzyme of FAO, carnitine palmitoyltransferase 1a (CPT1a) [**Figure 6B**]. We compared the kinetics of TIL-mediated killing of Mel526 (E:T = 3:1) and observed that etomoxir had insignificant impact on the killing mediated by TILs consistent with the expectation that NR derived T cells do not engage FAO (**Figure 6C**). By contrast, activating CPT1a with baicalin significantly increased the killing mediated by the TILs, confirming that the killing functionality can be augmented by promoting FAO (**Figure 6D**). The mechanism by which baicalin enables enhanced FAO is controversial, with reports suggesting either allosteric activation of CPT1a (Dai *et al*., 2018) or an increase in the mRNA levels of *CPT1a* (increased protein amount) (Chen *et al*., 2018). To investigate if baicalin increased the amount of CPT1a or was only an allosteric activator, we performed RT-qPCR on the MART-1 TIL harvested at 30 minutes, one, and two hours in HBSS with and without baicalin treatment. These results showed no significant differences in either *CPT1a*; or the fatty acid synthesis enzymes, fatty acid synthase (*FASN),* and Acetyl-CoA carboxylase 1 (*ACC1)* at any timepoint (**Figure 6G**). These data support the function of baicalin as an allosteric activator of CPT1a. In aggregate, these results demonstrate that modulating FAO can directly impact TIL survival and antitumor function targeting defined epitopes, at least *in vitro*.

**Figure 6.**
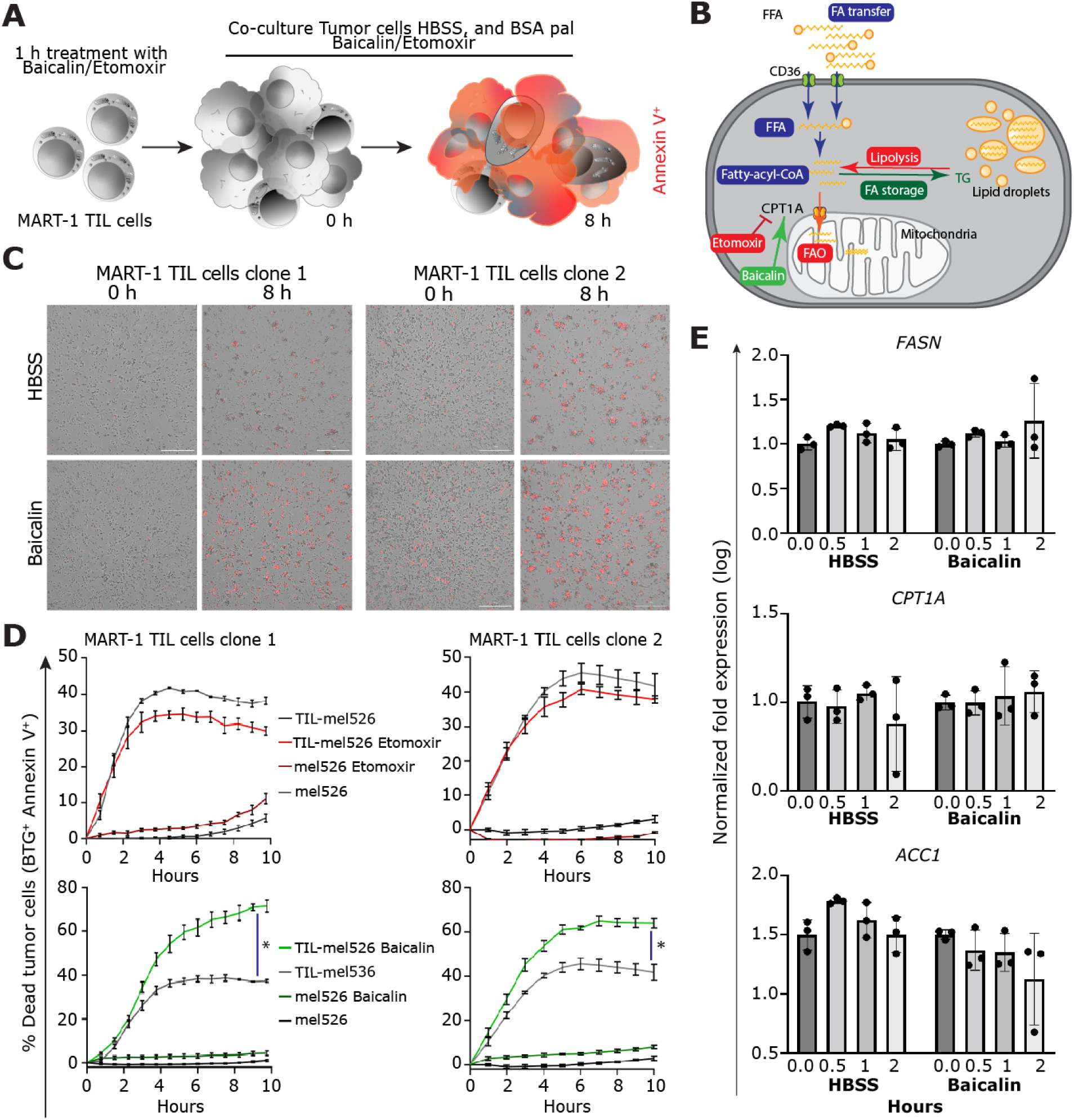
CPT1a stimulation increases the induction of target cell death by TILs. **(A)** Quantifying the cytolytic activity of TILs upon modulation of FAO. Clonal MART-1 specific TILs were pre-incubated for one hour in CM with baicalin (100 μM) or etomoxir (3μM). Target cells Mel526 were stained with biotracker green (BTG) and co-cultured with the TILs cells in HBSS and BSA-palmitoleic acid with or without either baicalin (100 μM) or etomoxir (3 μM); we added annexin V (AnnV) to the media to track the apoptosis. **(B)** The mechanism of action of baicalin and etomoxir within cells; both compounds target the rate-limiting enzyme of FAO, CPT1a. **(C)** Representative micrographs of the cell-mediated cytotoxicity between R TILs and NR TILs and their autologous tumor cells treated with baicalin 100 μM. **(D)** Comparison of the cell-mediated cytotoxicity between R TILs and NR TILs and their autologous tumor cells treated with (C) etomoxir 3 μM or (D) baicalin 100 μM. The cell-mediated cytotoxic assay was performed for 8 hours on an E:T ratio of 1:3; each condition was performed in triplicates. The percentage of dead cells was calculated as double-positive cells Biotracker-green (BTG)^+^annexin V (AnnV)^+^. The error bars denote the SEM. **(E)** Fold change in gene expression for TILs under starvation conditions (HBSS) with and without baicalin (100 μM). cDNA was prepared, and qRT-PCR was run to compare the *CPT1a*, *FASN*, and *ACC1* RNA levels. Relative gene expression quantification was normalized with *GAPDH* and *RPS18*. Experiments were performed in triplicates. Fold changes in gene expression analysis were performed using a Mann-Whitney test. *p* values for comparisons were computed using a non-parametric Mann-Whitney test. \**p* < 0.05, \*\**p* < 0.01,\*\*\*\**p* < 0.0001 in D and E.

**Figure 7.**
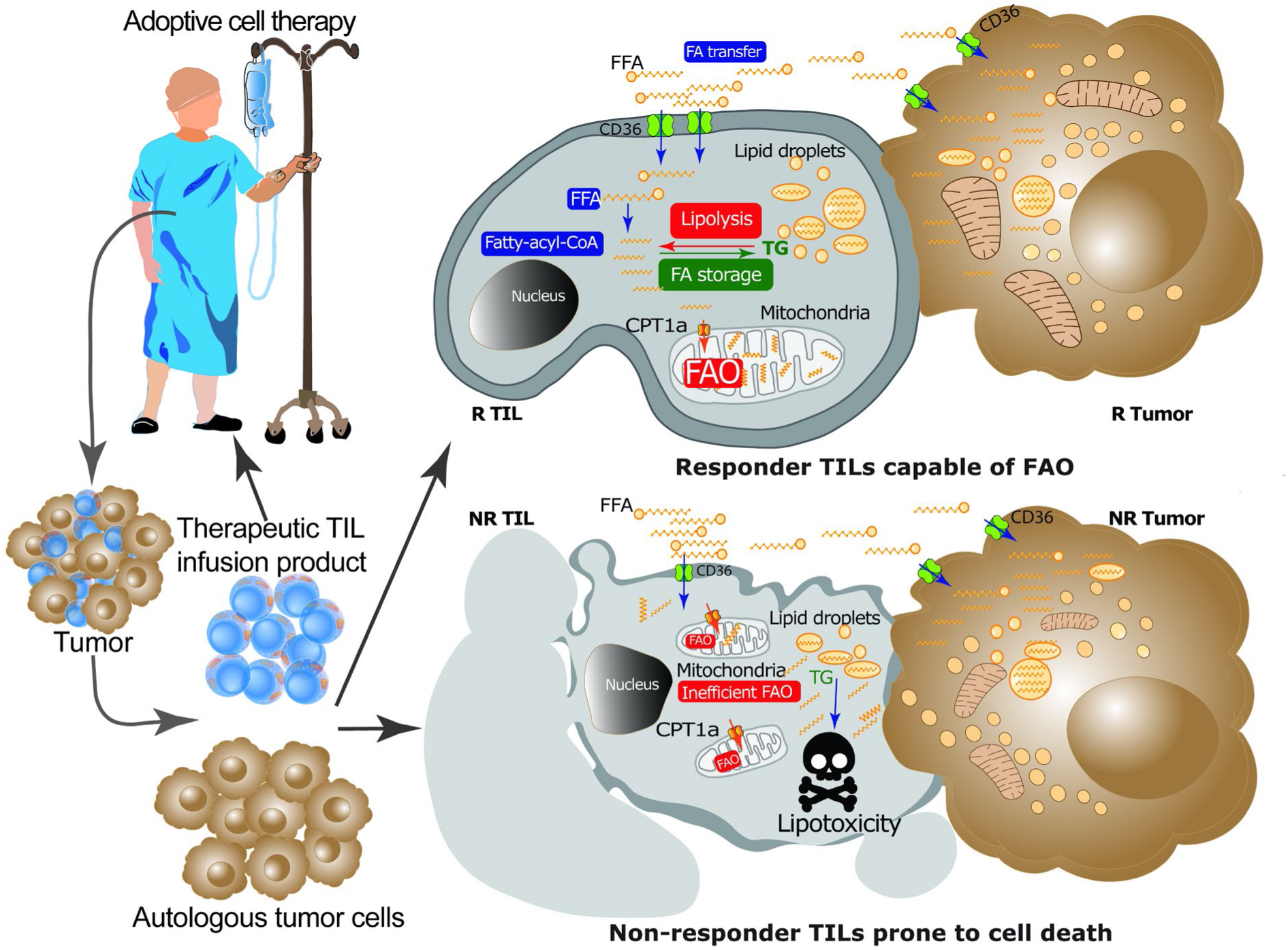
R TILs are capable of efficient FAO under conditions of nutrient limitation. Tumor cells from Rs and NRs are capable of efficient FA uptake and FAO under conditions of nutrient limitation. By contrast, although R and NR TILs can take up FAs, R TILs, co-cultured with autologous tumor cells under nutrient starvation, can perform efficient FAO. This allows the R TILs to survive lipotoxicity and preserve anti-tumor killing. NR TILs, however, cannot perform FAO and succumb to lipotoxicity, limiting their anti-tumor function. FFA = free fatty acid, TG triglycerides.

## DISCUSSION

Progressing tumors are highly evolved ecosystems that continuously adapt to create hostile microenvironments that enable the balance between supporting their own growth/dissemination and presenting a toxic environment to the antitumor immune cells. Successful immunotherapy should be able to address the challenges faced by the antitumor T-cells to enable a response. Since the function and survival of antitumor T-cells is thus context-dependent, we deployed a series of HABITAT assays that enable interaction between TILs being used for ACT and their autologous tumor cells to identify properties of TILs associated with clinical responses. Our results broadly illustrate that metabolic adaptation to utilize FAs by performing FAO is a feature of TILs associated with clinical response in ACT for metastatic melanoma.

FAs are essential to cancer cells since they enable membrane biosynthesis during rapid proliferation and allow metabolic plasticity under conditions of nutrient limitation (Vriens *et al*., 2019). The role of FAs during tumorigenesis and cancer progression is primarily through an elevated rate of *de novo* lipogenesis (Hoy *et al*., 2021; Vriens *et al*., 2019). *De novo* lipogenesis promotes the biosynthesis of saturated and monounsaturated FAs that, along with cholesterol, promote membrane rigidity (Rysman *et al*., 2010; Zhao *et al*., 2016). Despite this knowledge, and preclinical promise, translating the inhibition of lipogenesis by inhibiting FASN has largely yielded disappointing results (Falchook *et al*., 2021; Jones and Infante, 2015). In human melanomas (and other solid tumors), this can partially be explained by the fact that disseminated human melanomas are challenging to treat. As cancers metastasize, they undergo changes that enable their adaptation. From the perspective of the lipid composition of their membrane, cells that switch to single-cell migration (metastatic cells, EMT cells, amoeboid cells, circulating tumor cells [CTCs], etc.) harbor more unsaturated FAs and lower cholesterol. This switch in lipids enables greater membrane fluidity permitting cell deformability and promoting invasiveness and cellular migration. In our cohort, NR tumors harbor a greater frequency of migratory amoeboid cells. In human primary melanomas, amoeboid cells are enriched in the invasive front compared to the tumor body (Georgouli *et al*., 2019). Unlike cancer cells during tumorigenesis, metastatic migratory cells derive FAs through exogenous uptake. This uptake of FAs also confers a metabolic advantage since the cells can perform FAO, which is energetically more efficient than carbohydrate metabolism to generate ATP. Clinically: (1) the expression of CD36 (FA transporter) and FAO signatures is associated with metastatic progression and poor survival in multiple human tumors (Nath and Chan, 2016), and (2) higher baseline expression of FAO signatures or induction of FAO upon treatment is associated with treatment resistance in BRAF^V600E^-mutant melanomas (Aloia *et al*., 2019; Shen *et al*., 2020). Preclinically: (1) CD36^+^ cells are metastasis-initiating cells (Pascual *et al*., 2017), (2) lipid abundance is a hallmark of the TME in diverse tumors, including melanoma and pancreas (Manzo *et al*., 2020; Zhang *et al*., 2017), (3) tumor cells counteract increased lipid uptake and lipid peroxidation leading to ferroptosis by upregulating Glutathione peroxidase 4 (GPX4) that catalyzes the detoxification of lipid reactive oxygen species (Tsoi *et al*., 2018), and (4) promoting FAO through upregulation of either Acox1, Nur77 or YAP promotes survival of melanoma persister cells and metastasis (Lee *et al*., 2019; Li *et al*., 2018; Shen *et al*., 2020). Collectively, it is clear that lipid accumulation, uptake, and metabolism are necessary adaptations of metastatic melanomas leading to poor survival.

The accumulation of lipids and the concomitant glucose depletion within the TME present a metabolically hostile environment for T-cells. CD4^+^ T-cells, specifically regulatory T-cells (T_regs_), express CD36 to promote efficient lipid uptake and GPX4 to prevent ferroptosis (Xu *et al*., 2021a). Tregs’ immunosuppressive and pro-tumorigenic functions dampen antitumor immunity mediated by CD8^+^ T-cells. While quiescent and memory CD8^+^ T-cells perform FAO, activated CD8^+^ T-cells rely on glucose to fulfill their energy demands, and glucose deprivation directly impairs their antitumor functions, including the secretion of IFNγ (Manzo *et al*., 2020). In addition, the expression of PD1 inhibits glycolysis in CD8^+^ T-cells, promoting their functional exhaustion (Patsoukis *et al*., 2015). Lipids and FAs, on the other hand, directly dampen CD8^+^ T-cell function and survival. Clinical data on lipid uptake illustrates that: (1) CD8^+^ TILs in human melanomas, regardless of differentiation status, efficiently take up FAs (Zhang *et al*., 2017), (2) CD8^+^ TILs in pancreatic ductal adenocarcinomas (PDA) accumulate neutral lipids (Manzo *et al*., 2020), and (3) the survival of CD103^hi^ tissue-resident memory T-cells (T_RM_ cells), the likely dominant phenotype of TILs in gastric adenocarcinoma, was dependent on free FA (FFA) (Lin *et al*., 2020). Preclinical data from mice illustrate that: (1) CD8^+^ T_RM_ cells in the skin require exogenous lipid uptake and metabolism for their survival (Pan *et al*., 2017), (2) CD36^+^ CD8^+^ TILs in melanoma and colon cancer models are functionally exhausted, and undergo ferroptosis (Ma *et al*., 2021; Xu *et al*., 2021b), and (3) the accumulation of long-chain FA causes CD8^+^ TIL dysfunction in pancreatic cancers (Manzo *et al*., 2020). These data collectively illustrate that lipid uptake is a metabolic feature of CD8^+^ TILs and might be attributable to their tissue/tumor resident adaptation. However, uptaking lipids in the absence of compensatory mechanisms to utilize the lipids are detrimental to the function and survival of CD8^+^ TIL and can explain the exhaustion or absence (apoptosis) of CD8^+^ TIL in solid tumors.

Our data, derived from paired multi-omic profiling of the interactions between human TILs and their autologous tumor cells, enabled us to uncover an elegant metabolic adaptation that can facilitate TIL function in lipidic but otherwise nutrient-deficient conditions. In metabolic competition experiments that simulate nutrient competition found in the TME, tumor cells from both R and NR cells efficiently take up and utilize FAs. However, the critical difference is that CD8^+^ TILs from R samples of ACT efficiently perform FAO under conditions of metabolic competition, enabling their survival. By contrast, NR TILs are sensitive to lipotoxicity and undergo cell death under conditions of metabolic competition with autologous tumor cells. We did not observe significant differences in the expression of CD36 in the TILs of R and NR samples. This observation is consistent with preclinical data that showed no differences in FA uptake between CD36^+/+^ and CD36^-/-^ CD8^+^ TILs in melanoma or colon cancer models, suggesting redundancy in lipid transporters (Xu *et al*., 2021b). Our data also diverges from preclinical data from mice, illustrating that memory T-cells primarily derive lipids for FAO via FA synthesis from glucose (O’Sullivan *et al*., 2014). As described above, we submit that the human CD8^+^ TILs have fundamentally adapted for lipid uptake due to survival in a lipogenic TME. The ability of CD8^+^ TILs to engage FAO as a mechanism to preserve function and survival mimics the process used by metastatic tumor cells during metabolic adaptation. Engaging FAO to minimize lipotoxicity while supporting metabolism provides a pathway to fitter TILs for ACT compared to deletion of CD36 or overexpression of GPX4 protein that only will minimize lipotoxicity (Ma *et al*., 2021; Xu *et al*., 2021a). Preclinical data illustrate that systemic administration of fibrates (peroxisome proliferator-activated receptor [PPAR] agonists) can improve the antitumor function of OT-1 T-cells against the highly immunogenic B16-OVA model and synergize with anti-PD-L1 therapy in colon cancer models (Chowdhury *et al*., 2018; Zhang *et al*., 2017). The translatability of this approach is limited since systemic administration of drugs to increase FAO in TILs will also increase FAO in tumor cells, promoting their dissemination. This example highlights one of the central challenges in translating therapeutics-targeting metabolism; since the metabolic pathways that enable plasticity and adaptation to harsh microenvironments are shared between metastatic tumor cells and the antitumor immune cells, systemic administration will always provide a combined (but weighted) integration of the impact on both sets of cells. In this context, ACT is unique since there is an opportunity for *ex vivo* genetic manipulation of the T-cells (like the enforced expression of the peroxisome proliferator-activated receptor-gamma coactivator one alpha [PGC1α] protein) before the infusion to enable their sustained survival and function (Dumauthioz *et al*., 2021).

Beyond ACT, T-cell adaptation to lipid-rich environments is also significant in the context of ICB. For example, profiling plasma metabolites showed that acylcarnitines were elevated in NR compared to R cells upon anti-PD1 treatment in lung cancers (Hatae *et al*., 2020). Since activation of FAO promotes intracellular transport of acylcarnitines, leading to a concomitant decrease in plasma, this observation suggests that NR immune cells do not efficiently engage FAO. Furthermore, on-treatment profiling of metastatic melanoma patients treated with anti-PD1 showed that upregulation of FA metabolism upon treatment is associated with clinical responses across multiple patient cohorts (Du *et al*., 2021). Lastly, computational discovery of tumor-resilient T-cells (Tres) properties that can predict responses to ICB or ACT in diverse tumors, including metastatic melanoma, identified the fibroblast growth factor intracellular-binding (FIBP) protein, a cholesterol modulator, as the top negative regulator of Tres (Zhang *et al*., 2022).

In summary, engagement of FAO in T-cells under conditions of metabolic competition and nutrient starvation represents an adaptation that promotes their function and survival. Furthermore, selective engagement of FAO in T-cells provides a pathway to clinical responses in cancer immunotherapy.

## METHODS

### Patient selection

Metastatic melanoma tumor and TILs were expanded from surgical resection tissue under the protocol (2004–0069) approved by the Institutional Review Board (IRB) of the University of Texas MD Anderson Cancer Center (Houston, TX) and an FDA-approved Investigational New Drug (IND) application (NCT00338377). As the Declaration of Helsinki describes, the study was conducted in compliance with Good Clinical Practice concerning human medical research.

For this study, we used 16 samples (eight responders and eight nonresponders with known overall survival). All patients with stage III and IV melanoma were over 18 years old. The characteristics of patients are described in **Supplementary Table 3**. The methodology to expand TIL has been previously described (Forget *et al*., 2018; Radvanyi *et al*., 2010). Briefly, TILs were initially expanded from the harvested surgical tumor (pre-REP), followed by a rapid expansion protocol (REP). The TILs collected were infused into treated patients following a pre-infusion lymphodepletion chemotherapy regimen and along with high doses (HD) of IL-2 therapy (Proleukin^TM^; Prometheus, San Diego, CA) [**Figure S1**]. In addition, the patient’s peripheral blood lymphocyte (PBL) samples were obtained weekly during follow-up visits and processed by the MDACC Melcore Lab. Infusion products and PBL samples were cryopreserved and stored in liquid nitrogen. Samples are referred to by their TIL number displayed in **Table 2**. The HLA-A locus of the samples was typed at the MD Anderson H.L.A. Typing Laboratory and shown in **Table 3**.

**Table 3.**
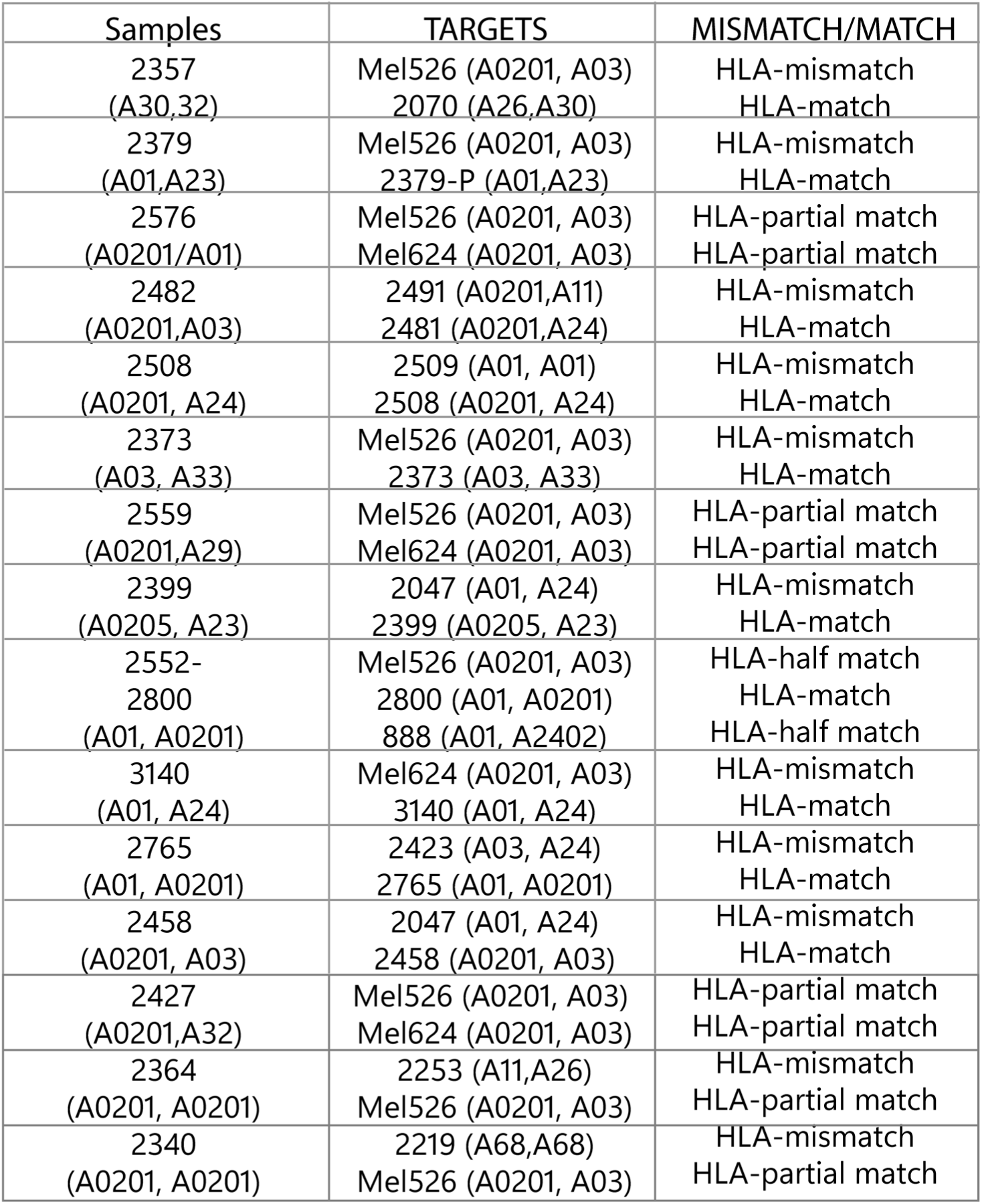
List of the human leukocyte antigen (HLA) alleles present on the TILs of the sample used in this study.

| Samples | TARGETS | MISMATCH/MATCH |
| --- | --- | --- |
| 2357<br>(A30,32) | Mel526 (A0201, A03)<br>2070 (A26,A30) | HLA-mismatch<br>HLA-match |
| 2379<br>(A01,A23) | Mel526 (A0201, A03)<br>2379-P (A01,A23) | HLA-mismatch<br>HLA-match |
| 2576<br>(A0201/A01) | Mel526 (A0201, A03)<br>Mel624 (A0201, A03) | HLA-partial match<br>HLA-partial match |
| 2482<br>(A0201,A03) | 2491 (A0201,A11)<br>2481 (A0201,A24) | HLA-mismatch<br>HLA-match |
| 2508<br>(A0201, A24) | 2509 (A01, A01)<br>2508 (A0201, A24) | HLA-mismatch<br>HLA-match |
| 2373<br>(A03, A33) | Mel526 (A0201, A03)<br>2373 (A03, A33) | HLA-mismatch<br>HLA-match |
| 2559<br>(A0201,A29) | Mel526 (A0201, A03)<br>Mel624 (A0201, A03) | HLA-partial match<br>HLA-partial match |
| 2399<br>(A0205, A23) | 2047 (A01, A24)<br>2399 (A0205, A23) | HLA-mismatch<br>HLA-match |
| 2552-2800<br>(A01, A0201) | Mel526 (A0201, A03)<br>2800 (A01, A0201)<br>888 (A01, A2402) | HLA-half match<br>HLA-match<br>HLA-half match |
| 3140<br>(A01, A24) | Mel624 (A0201, A03)<br>3140 (A01, A24) | HLA-mismatch<br>HLA-match |
| 2765<br>(A01, A0201) | 2423 (A03, A24)<br>2765 (A01, A0201) | HLA-mismatch<br>HLA-match |
| 2458<br>(A0201, A03) | 2047 (A01, A24)<br>2458 (A0201, A03) | HLA-mismatch<br>HLA-match |
| 2427<br>(A0201,A32) | Mel526 (A0201, A03)<br>Mel624 (A0201, A03) | HLA-partial match<br>HLA-partial match |
| 2364<br>(A0201, A0201) | 2253 (A11,A26)<br>Mel526 (A0201, A03) | HLA-mismatch<br>HLA-partial match |
| 2340<br>(A0201, A0201) | 2219 (A68,A68)<br>Mel526 (A0201, A03) | HLA-mismatch<br>HLA-partial match |

### IFNγ ELISA

Successfully expanded pre-REP TILs were tested for anti-tumor reactivity (autologous setting) by measuring IFNγ production in the supernatant of a co-culture of TIL and autologous tumor cells as previously described (Radvanyi *et al*., 2012). In some cases, a semi-allogenic melanoma cell line with at least one matched HLA-A allele was used as a target (**Table 3**). IFNγ ELISA kits were obtained from Pierce^TM^.

### Generation of autologous tumor

Melanoma cells were generated as previously described (Andrews, Oba, *et al*., Nature Communications 2022). Briefly, each specimen from a metastatic melanoma tumor was collected and processed into a single cell suspension by incubation with an enzymatic digestion cocktail (0.375% collagenase type I, 75 µg/mL hyaluronidase, and 250 U/mL DNase-I) in tumor digestion medium (RPMI-1640 containing 10 mM HEPES, 1% penicillin/streptomycin and 20 μg/mL gentamicin; Gibco/Invitrogen) in a humidified incubator at 37°C with 5% CO2 with gentle rotation for 2-3h. The tumor digest was filtered through a 70 µm filter, washed in sterile PBS, and re-suspended in serum-free media, which was then plated in one well of a 6-well culture plate and incubated at 37°C in a 5% CO_2_ atmosphere. After 24 hours, the media was replaced with fresh complete tumor media, comprised of RPMI-1640 supplemented with 1% GlutaMAX, 10% FBS, 1% penicillin/streptomycin, 20 μg/mL gentamicin, 50 μM β-mercaptoethanol (Gibco/Invitrogen), 10 mM HEPES, and 5 μg/mL insulin-selenium-transferrin (Gibco/Invitrogen). Cells were grown-on in enriched DMEM/F12 culture media (Gibco/Invitrogen) supplemented as above and with 1 mM sodium pyruvate and MycoZap-PR (Lonza). Cultures were deemed established when the cells stained positive for a melanoma tumor marker (MCSP-1 Miltenyi, cat# 130-117-347) and negative for a fibroblast marker (CD90, BD Pharmingen, cat# 561558). Tissues were harvested primarily by surgical resection, and cell lines were generated by the MD Anderson Cancer Center TIL Laboratory between 2007 and 2014. All MDACC cell lines were used within ten passages of tumor line establishment; cells were cryopreserved and kept in stocks in liquid nitrogen until use.

### Tumor cell lines and co-culture

All the tumor cells used in the paper tested negative for mycoplasma (MycoAlert^®^, Lonza and MycoSensor qPCR^®^, Agilent). The metastatic melanoma cells were cultured in RPMI 1640 (Corning) supplemented with 10% FBS (Atlanta), 1 x L-glutamine (Corning), 1 x HEPES (Gibco/Invitrogen), 1 x Pen-Strep (Corning), 1 % Insulin Transferrin and Selenium (Gibco/Invitrogen), 20 mg/ml Gentamicin (Gibco/Invitrogen). TILs were grown in TIL culture media as follow RPMI 1640 (Corning) supplemented with 10% human sera (AB), 1x L-glutamine (Corning), 1x HEPES(Gibco/Invitrogen), 1x Pen-Strep (Corning) 1 mmol/L Sodium pyruvate (Corning) with 6000 IU/ml IL2 (hrIL-2) (Novartis) in 24-well plates. Melanoma A526 tumor cell line (ATCC) was cultured in the same media as the HLA-matched metastatic melanoma lines.

Co-culture TILs-melanoma tumor cells were done on a ratio of 5:1 or 3:1, using TIL culture media or in starvation conditions in HBSS (Gibco) for 6, 12, or 24 hours as described in each experiment.

### TCR Vβ sequencing and tracking

Total DNA was isolated from sorted TIL and PBMCs using the Qiagen AllPrep Kit according to the manufacturer’s instructions. TCRβ CDR3 sequencing was performed using the Adaptive Biotechnology (Seattle, WA) pipeline. Survey level sequencing was performed on DNA extracted from TIL, and deep level sequencing was performed on DNA extracted from PBMC. As previously described, Clonality was determined using Immunoseq (Robins *et al*., 2009). Tracking data was extracted from Immunoseq and analyzed at MD Anderson.

### Reagents

2NBDG (2-(N-(7-Nitrobenz-2-oxa-1,3-diazol-4-yl)amino)-2-deoxyglucose), Cell Trace^TM^ Far Red, MitoTracker® Green, MitoTracker® Red CMXRos, MitoTracker® Deep Red FM, Bodipy^TM^ 558/568 (C_12_), and Bodipy^TM^ 493/503 were purchased from Invitrogen. Live and dead fixable Aqua dead cell staining Aqua (Invitrogen) and Ghost Dye^TM^ Violet 450 were from Tonbo Bioscience. IL-2 was from Proleukin^TM^. Baicalin and etomoxir (purity≥98%) were purchased from SelleckChem. BSA-palmitate was purchased from Cayman Chemical Company. For cell experiments, compounds were dissolved in dimethyl sulfoxide (DMSO), and the final concentration of DMSO did not exceed 0.1%.

### PDMS nanowell array fabrication and TIMING

The in-house fabrication of the PDMS nanowell array was previously described (Liadi *et al*., 2015). Timelapse Imaging Microscopy In Nanowell Grids (TIMING) method allows a high throughput time-lapse and single-cell level imaging of thousands of nanowells containing 1-4 cells. Effector (TILs) and target cells (autologous tumor cell lines) were labeled respectively with 1μM PKH67 and PKH26 fluorescent dyes (Sigma-Aldrich) according to the manufacturer’s protocol and loaded on the chip array. The array was immersed in phenol red-free cell-culture media containing a dilution of 1:60 Annexin V Alexa Fluor 647 (AF647) (Life Technologies) to detect cell apoptosis. Arrays were imaged for 6-8 hrs at an interval of 5 minutes using an Axio fluorescent microscope (Carl Zeiss) utilizing a 20x 0.45 NA objective, a scientific CMOS camera (Orca Flash 4.0), a humidity / CO2 controlled chamber, and the tile function of the Zen software.

### Flow cytometry

Antibodies against human CD3 FITC, CD4 PerCP Cy 5.5, CD8 APC Cy 7, CD69 APC, and CD28 PE Cy 7 were purchased from BD Biosciences. Antibody against human CD279 (PD1 clone EH12.2H7, PECy 7), CD3 PerCP-Cy5.5, CD8 PE, and FITC were obtained from Biolegend.

TILs were co-cultured with their respective autologous tumor cell for 12 hours, and then TILs were stained with surface antibodies CD3^+^, CD8^+^, and CD69^+^ cells and were sorted for RNA-seq. Cell sorting was performed on the FACS Aria Fusion Cell Sorter (BD Bioscience). Unstained and single-stained cells were used as controls. Dead cells were excluded using Aqua live/dead staining.

For the immune metabolism experiments, pre-stained tumor cells with Cell Trace Far Red (Thermo Fisher Scientific) and the corresponding TILs were washed and incubated with prewarmed media with 100 μM of MitoTracker Green (Thermo Fisher Scientific), 100 μM of MitoTracker Red CMXRos (Thermo Fisher Scientific) for 20 minutes at 37 °C, or 1 μM Bodipy 558/568 (C_12_) [Thermo Fisher Scientific] for 12 hours at 37 °C in a six-well plate. Cells were washed and stained with antibodies for cell surface antigens CD3, CD4, and CD8. For analysis of lipid content, 1 μg/ml Bodipy 493/503 was added in a 1x Permeabilization Buffer for one hour at four °C. Cells were analyzed on the LSR. Fortessa X-20 analyzer (BD Biosciences). Unstained and single-stained cells were used as controls. Dead cells were excluded using Ghost live/dead staining Violet 450 (Tonbo bioscience). The samples were read LSR. Fortessa X-20 analyzer (BD Biosciences). Furthermore, data analysis was conducted using FlowJo 10.8.1 software (Tree Star, Inc.).

### RNA sequencing

Total RNA from one hundred thousand TILs was extracted using a commercially available RNA isolation kit (Macherey-Nagel) following the manufacturer’s recommendations and, later, treated with DNase to remove DNA contamination. RNA quantity and quality were confirmed with the Qubit (Thermo Fisher Scientific) and the Agilent 2100 Bioanalyzer. A total of 5 µg of RNA with a minimal RNA Integrity Number (RIN) of 7.0 was used to prepare the cDNA libraries with the commercially available SMARTer Stranded Total R.N.A. Seq Kit (Takara), as per the manufacturer’s protocols. The libraries were pooled, and a 76 bp paired-end sequencing was performed on an Illumina NextSeq 500 platform using a high output flow cell to yield a minimum of 10 million reads per library (range 4–8 million). The melanoma tumor RNA sequencing library and sequencing were performed as previously described (Andrews *et al*., 2022).

### Bioinformatic analysis

Firstly, fifteen bases were trimmed from the 3’ and 5’ sites in all reads to remove the adapter sequences and eliminate the nonuniform distribution caused by the biased selection of reads or contamination of other sequences. Next, the RNA-seq reads were aligned with the HISAT2 (2.0.5) program from Johns Hopkins University (Kim *et al*., 2015) and mapped to the human reference genome (GRCh37/hg19). Later, differentially expressed genes (DEGs) were identified using the DESeq2 tool that tests for differential gene expression based on a model using the negative binomial distribution. Finally, the differentially expressed pathways were identified by the “GAGE” package using the gene sets downloaded from the “Molecular Signature Database” of the Broad Institute and also by using the Gene Set Enrichment Analysis (GSEA) software (Subramanian, 2005), Cytoscape open-source software (Shannon *et al*., 2003).

### Mass spectrometry

A total of one million autologous tumor cells (3 R and 3 NR) were lysed using a lysis buffer containing 6 M urea and 2 M thiourea in 50 mM ammonium bicarbonate. This was done in two technical replicates. Protein concentrations were determined using the Bradford assay. A total of 50 µg of protein was in-solution digested as follows: Lysates were reduced with one mM DTT, alkylated using five mM iodoacetamide, diluted 1:4 with 50 mM ammonium bicarbonate, and digested over-night with Lys-C-Trypsin mix (1:100 enzyme to protein ratio) and trypsin (Promega; 1:50 enzyme to protein ratio). Following the digestion step, the peptides were acidified using 1% trifluoroacetic acid, separated using strong cation exchange (SCX) chromatography in a stage tip format, and purified on C-18 (3M) stage tips (Rappsilber *et al*., 2003).

### LC-MS/MS data acquisition and processing

LC-MS/MS runs were performed on the EASY-nLC1000 UHPLC (Thermo Scientific) coupled to the Q-Exactive HF mass spectrometer (Thermo Scientific) (Scheltema *et al*., 2014). Peptides were separated using a 50 cm EASY-spray PepMap column (Thermo Scientific) with a water-acetonitrile gradient of 140 minutes for each fraction. Buffer A was 0.1% formic acid; buffer B was 80% acetonitrile and 0.1 % formic acid. The resolutions of the MS and MS/MS spectra were 60,000 and 30,000, respectively. MS data were acquired in a data-dependent mode with target values of 3,000,000, and 500 for MS and MS/MS scans, respectively, using a top-10 method.

Raw MS data were processed using MaxQuant (Cox and Mann, 2008; Tyanova *et al*., 2016). Database search was performed with the Andromeda search engine based on the human Uniprot database. The settings included carbamidomethyl cysteine as a fixed modification, methionine oxidation, and N-terminal acetylation as variable modifications. The forward/decoy approach was used to determine the false discovery rate (FDR) and filter the data with a threshold of 1% FDR for both the peptide and the protein levels. In addition, the “match between runs” option was enabled.

### Proteomics data analysis

The proteomics data were analyzed using Perseus (Tyanova and Cox, 2018). A total of 9362 protein groups were identified and quantified using the label-free quantification (LFQ) algorithm (Cox *et al*., 2014). Reverse proteins, proteins that were only identified by site, and potential contaminants (except for keratins) were filtered out. The data were log2-transformed, and the protein groups were filtered to keep those with a valid value in at least five samples in one group, reaching a final list of 7087 protein groups. A Student’s t-test was done to get the differentially expressed proteins between the Rs and the NRs cell lines (FDR < 0.05). In order to determine the biological processes enriched in the proteomics and the RNA-seq data, 2-D enrichment analysis (Cox and Mann, 2012) was done on the t-test difference between R and NR in the proteomics and the RNA-seq data (FDR < 0.02).

### Genome-Scale Metabolic Model

Transcriptomic data was obtained from tumor cells and TILs, and proteomic data were obtained from tumor cells. Samples were labeled as Rs or NRs for each cell type. Differential expression (DE) analysis was performed on the transcriptomic data within each cell type to evaluate the difference between NR and R samples. The results of the DE analysis (expression fold-change and significance of each gene) were subsequently analyzed with gene set analysis, using either the Molecular Signatures Database (MSigDB) hallmark gene set collection, or the gene-metabolite interaction network extracted from the Human1 model. Transcriptomic and proteomic data were integrated with Human1 to extract a GEM specific to each sample. The network structures of the resulting sample-specific GEMs were compared to identify differences in reaction or pathway content between Rs and NRs in TIL and tumor cells.

### Confocal analysis

TILs were washed and incubated for 12 hours with oleic acid (Sigma-Aldrich) or one mM of Bodipy 568/550 (C_12_) [Thermo Fisher Scientific]. In addition, cells were washed and stained with prewarmed media with 100 μM of MitoTracker Green (Invitrogen), or 100 μM of MitoTracker Far Red (Invitrogen), Hoechst 33258 (Invitrogen), with/without 1 μg/ml Bodipy 493/503 (Invitrogen) for 20 minutes at 37 °C. Cells were washed and seeded in a 96-well dark plate (coverslip bottom) with their respective autologous tumor cell line. Cells were analyzed on a Nikon spinning disk confocal analyzer at 0 and 24 hours.

To quantify the confocal images, we extracted the TIFF images for lipid droplets (Bodipy 493/503), mitochondria, and fatty acids (Bodipy 558/568 [C_12_] and processed them in ImageJ (National Institutes of Health [NIH], USA). We cropped cells for 2D analysis of lipid droplets to ensure the detected lipid droplets belonged to a single cell. Then, we binarized the images for lipid droplets, and using analyze particle plugin; we calculated the area of lipid droplets in single cells.

To analyze the 3D images of mitochondria and fatty acids, we used a series of plugins, including 3D watershed, 3D object counter plugin (Bolte and Cordelières, 2006), and 3D ROI Manager plugin (Ollion *et al*., 2013), to measure the pixel values and total volume of each mitochondrion and fatty acid in the cells. Then, we imported these data into R (v 4.0.2) and calculated the total volume of fatty acid per cell and Pearson’s coefficient of fatty acid overlapping with mitochondria per cell using the formula:

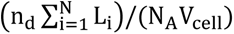

R and G stand for pixel values on fatty acid image and mitochondria image per cell, respectively. We used Graph Pad Prism software for statistical testing.

### Cell-mediated cytotoxic assay

Mel-526 cells stained with BioTracker 490 green cytoplasmic membrane fluorescent dye (Millipore Sigma) were seeded on poly-L Lysine pretreated 96 well plates. Pretreated effector cells with the inhibitor etomoxir and the baicalin per one hour were added to the corresponding wells at an E:T = 3:1 density in HBSS media, supplemented with 50 IU/ml of IL-2, annexin V Alexa Fluor 647 (AF647) (Life Technologies), BSA-palmitate (Cayman Chemical), Etomoxir (SelleckChem), Bailcain (SelleckChem) in the corresponding wells. The cells were tracked every 45 minutes for the next ten hours using Cytation 7 (Byotek). The killing assay was analyzed using double mask detection (Biotracker green positive cells and Annexin V positive cells).

### RNA isolation, cDNA preparation, and qRT-PCR

The clonal TIL line that recognizes MART-1 antigen was treated with baicalin for the indicated time and later lysed in RNeasy lysis buffer (RLT) from the RNeasy mini kit (Qiagen) to extract RNA. Next, 1 μg of total RNA was converted to cDNA using the High-Capacity cDNA Reverse Transcription kit (Invitrogen). Real-time PCR was performed with primers for *CPT1a*, *FASN*, *ACC1*, *Actin*, and *RPS18* listed in **Table 5**, using the SsoFast^TM^EvaGreen^®^ supermix with low ROX (BIORAD) on AriaMx Real-time PCR system (Agilent Technologies). Relative gene expression quantification was normalized with Actin and RPS18, and mRNA expression was calculated using the 2 ^-ΔΔCt^ methods. Analysis was carried out in triplicates.

**Table 4.**
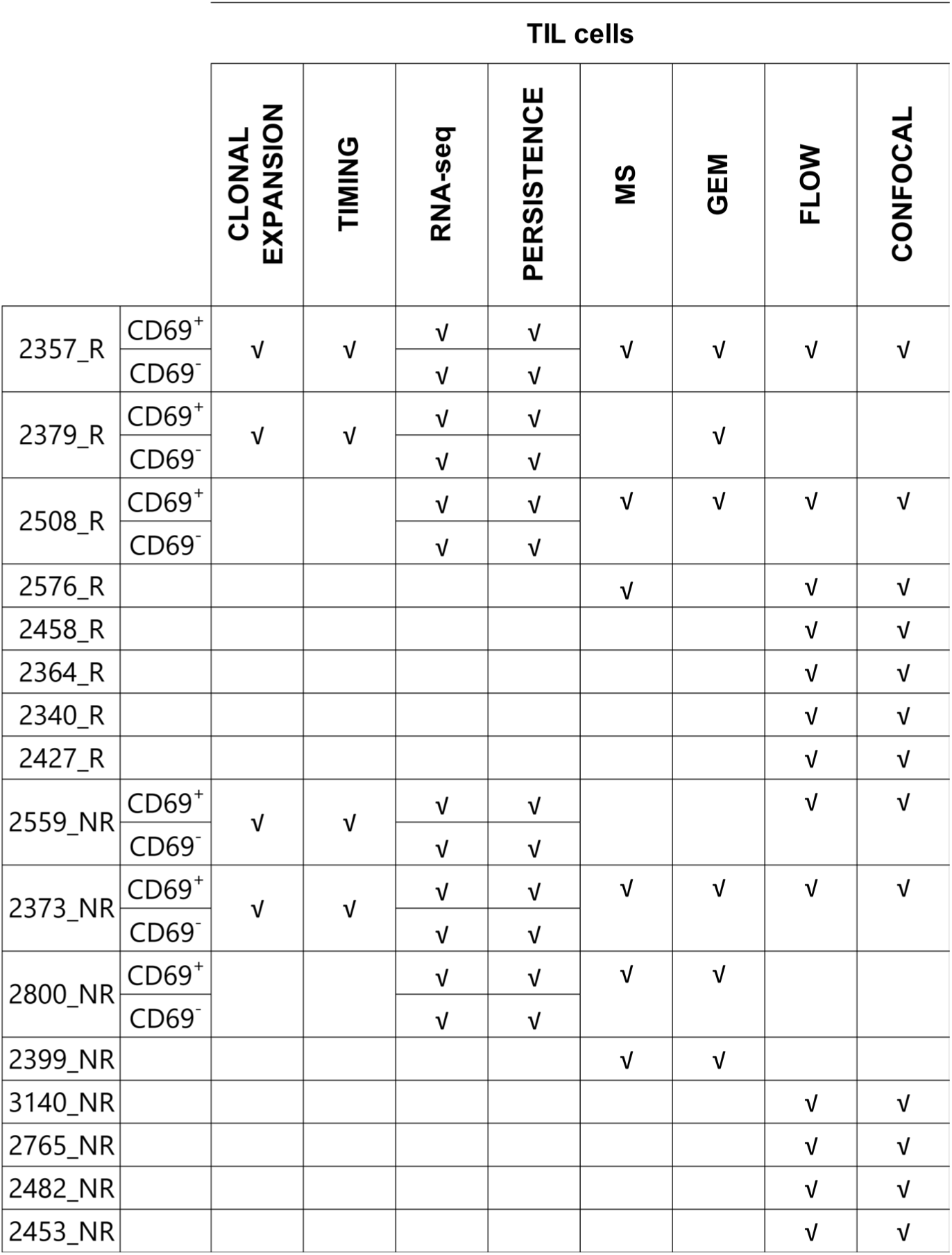
List of samples used in each experiment

**Table 5.**
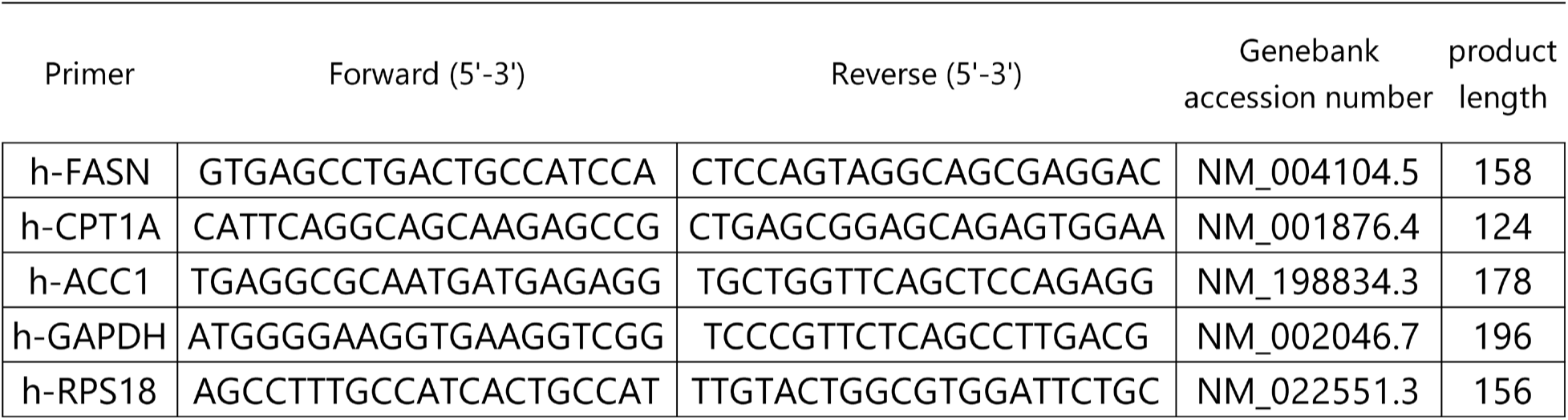
Primers used for qRT-PCR analysis of TILs.

### Statistical analysis

The data were processed using GraphPad Prism 8.0 software (CA, USA). As appropriate, differences between groups or experimental conditions were determined using nonparametric Mann-Whitney tests. The graphs show the mean value. Spearman correlation analyses were used as indicated. Two-sided P-values <0.05 were considered statistically significant and in the figures are indicated as *, *p* < 0.05; **, *p* < 0.01; ***, *p* < 0.001, and ****, *p* < 0.0001.

### Author Contributions

NV, PH, CH, CB, and MMP designed the study. XA & AR fabricated PDMS nanowell arrays. CH and MMP performed the sorting. MMP performed the TIMING and TILs RNAseq. SW performed RNAseq of tumor cells. MMP and AA analyzed the TIMING data. XA, MF, and MMP performed flow cytometry. TG & MH performed and analyzed the mass spectrometry. JR performed the genome-scale metabolic modeling. MF and JTA analyzed single-cell RNA-seq. MF performed image segmentation and analysis of confocal images. MMP and MK designed the figures. CB, CH, and NV supervised the study.

## Acknowledgments

We would like to acknowledge the MDACC Flow Cytometry Core Facility for the FACS sorting and analyzers (NCI P30CA16672). We thank Intel for the donation of a cluster.

---

**Supplementary figure 1.**
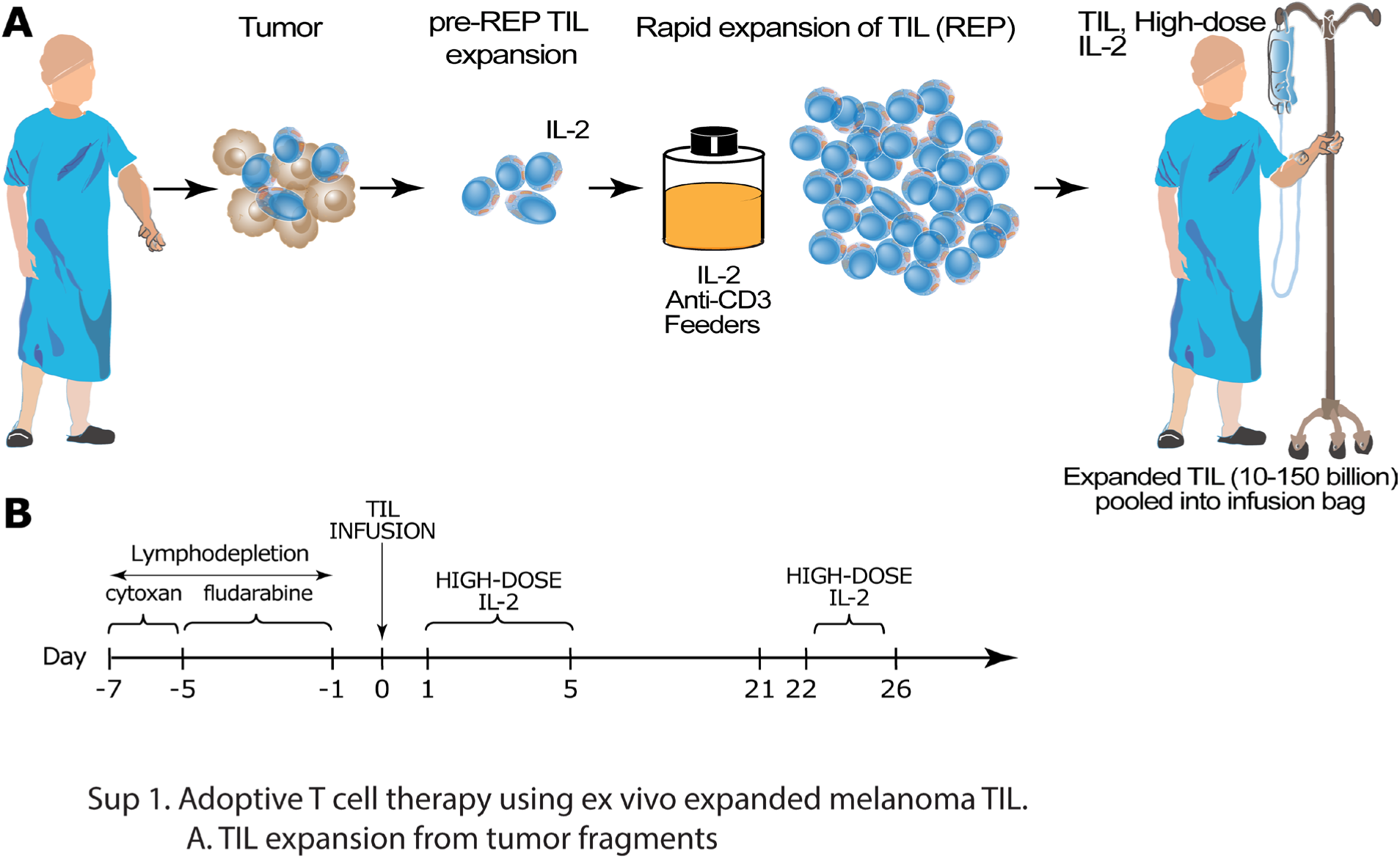
Adoptive T-cell therapy using *ex vivo* expanded melanoma TILs. **(A)** TILs are harvested from tumors, expanded, and infused into the patient with high doses of IL-2. **(B)** Clinical timeline for the treatment of patients using ACT.

**Supplementary figure 2.**
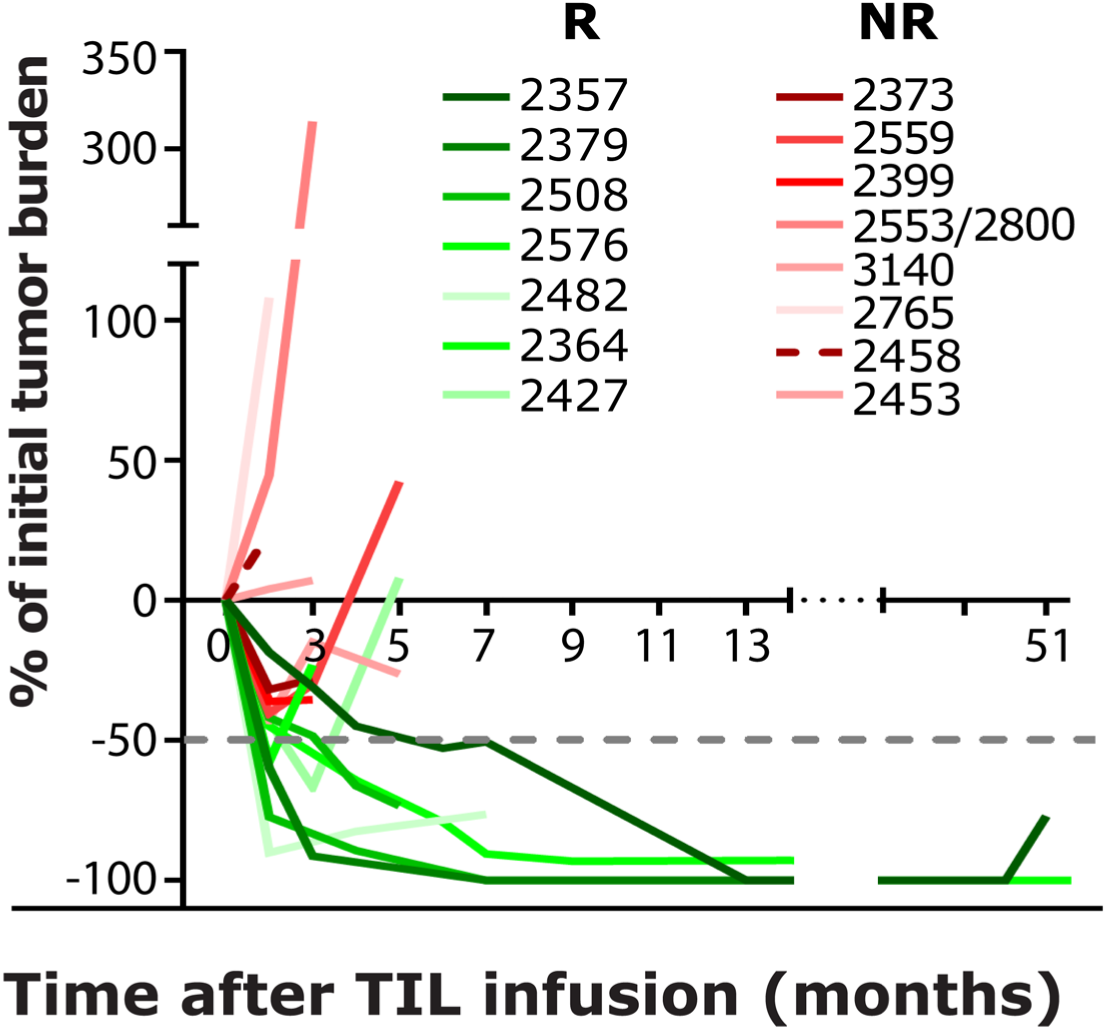
Longitudinal differences in tumor burden between R and NR samples after infusion of TILs. Spider plot showing the tumor regression through time after infusion of TILs (green, R, and in red, NR samples).

**Supplementary figure 3.**
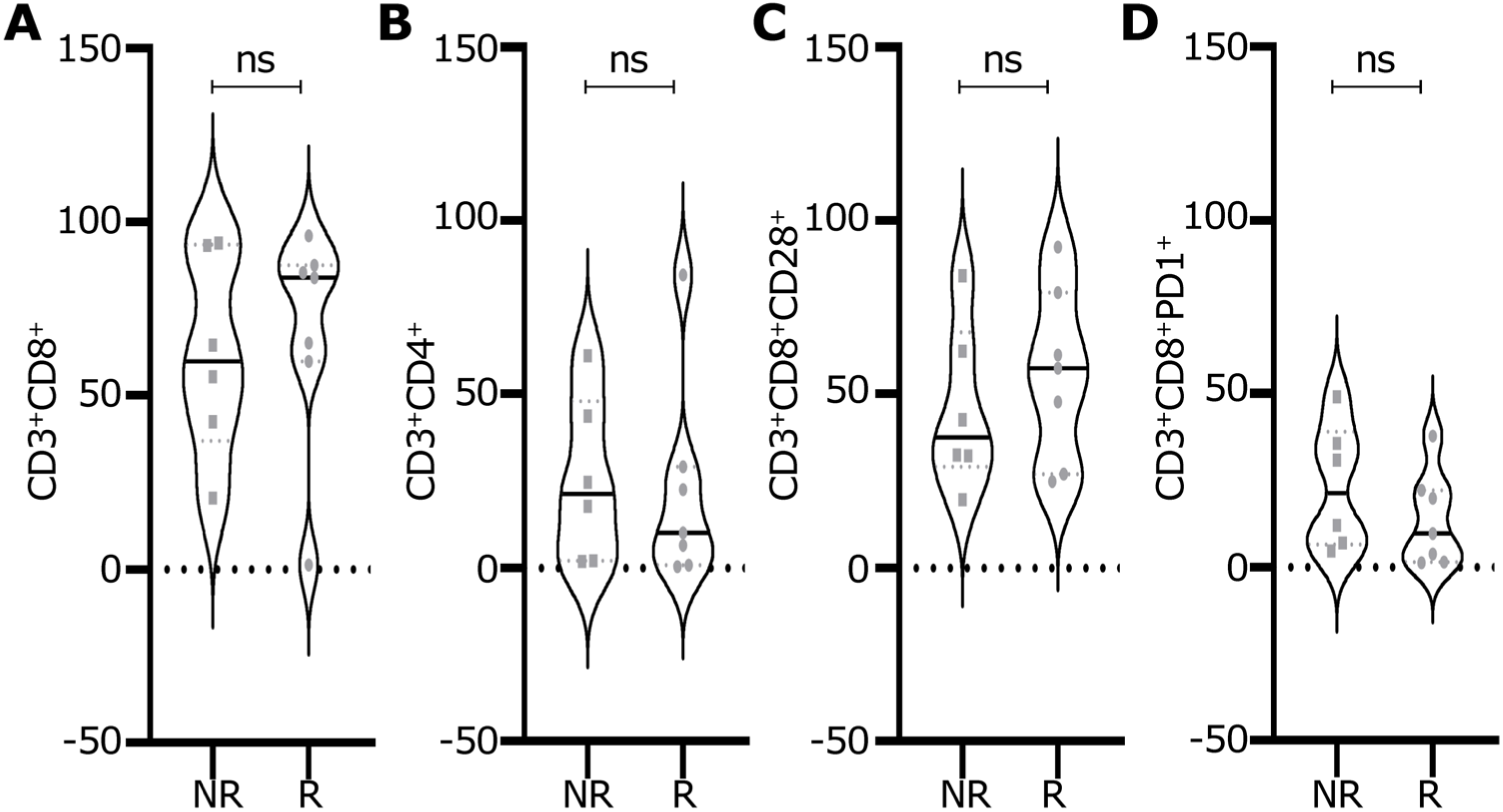
TIL Infusion products comprising R and NR patients are phenotypically similar. Infusion product TILs from Rs and NRs were stained with fluorescent antibodies to evaluate the percentage of surface markers by flow cytometry. (A-D) frequencies of the appropriate staining are indicated. (n = 12, six R, six NR TILs) Samples were analyzed by flow cytometry. *p* values for comparisons were computed using a non-parametric t-test. ns = no significance.

**Supplementary figure 4.**
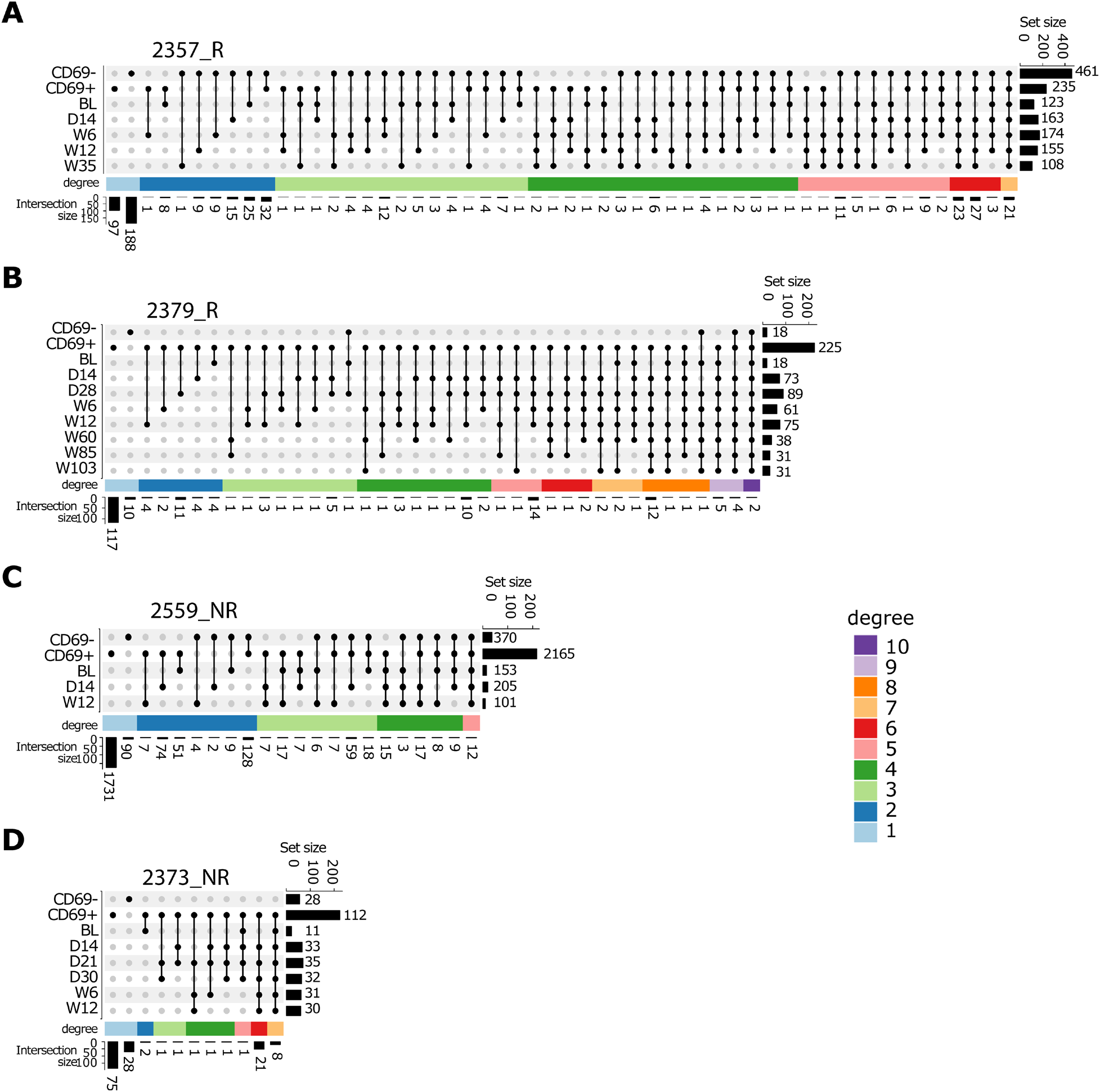
R and NR TILs are composed of diverse clonal populations that expand after infusion. TIL from R and NR patients’ samples were co-incubated with autologous tumor cells. Flow-sorted CD8^+^CD69^+^ (activated) and CD8^+^CD69^-^ (unactivated) T-cells were used to determine the TCRβ CDR3 sequences by high-throughput sequencing. The persistence of the cells over time is visualized as an UpSet plot (Ewing, 2021; Gu, 2022; Gu *et al*., 2016; Neuwirth, 2022; Team, 2021; Wickham and Bryan, 2022). Each dark circle represents the first timepoint at which a set of clones are detected. The connected lines and connected dark circles track the evolution of the persistence of that particular set of clones. For the 2357_R sample, the first two circles represent the number of CD69+CD8+ and CD69-CD8+ clones detected only in the infusion product but never after that. The bar graph on the right represents the total number of unique clones in different sets (set size), which are CD69^-^CD8^+^ and CD69^+^CD8^+^ in the infusion product, as well as the unique clones from the patient at different time points: BL = Baseline, D = day, W = week (top right). The column graph at the bottom represents the number of unique clones common across the infusion product and the patient’s clonal diversity across time (intersection size). Two R TIL samples (**A**-**B**) and two NR TIL samples (**C**-**D**) are shown.

**Supplementary figure 5.**
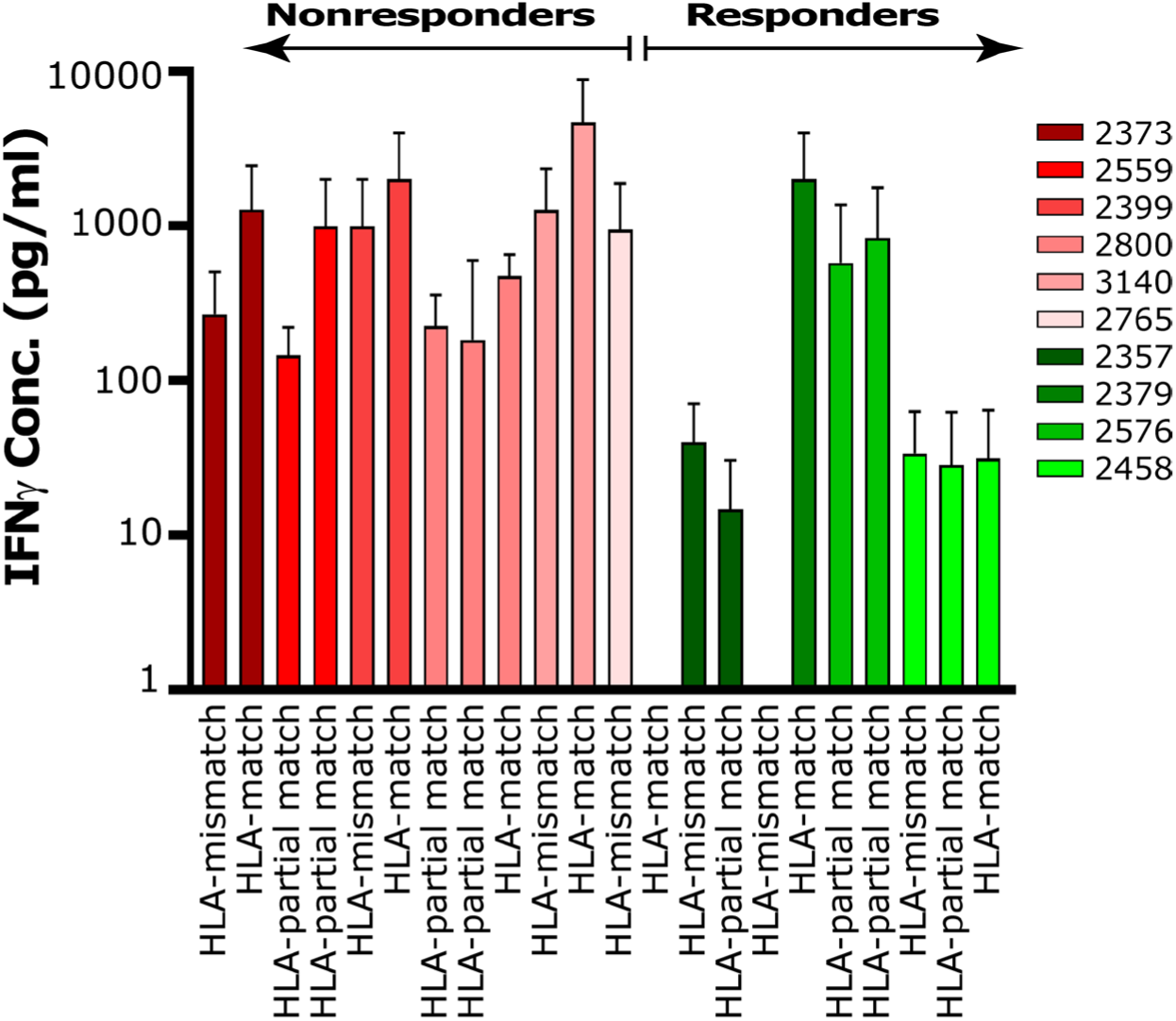
TIL infusion products from R and NR samples showed reactivity to the HLA-matched tumors. TILs were co-cultured with either HLA matched or mismatched tumors at a 1:1 ratio of 1 x 10^5^ density of cells in a 96-well plate for 24 h. The co-culture supernatants were tested using ELISA (supernatant was collected from triplicates).

**Supplementary figure 6.**
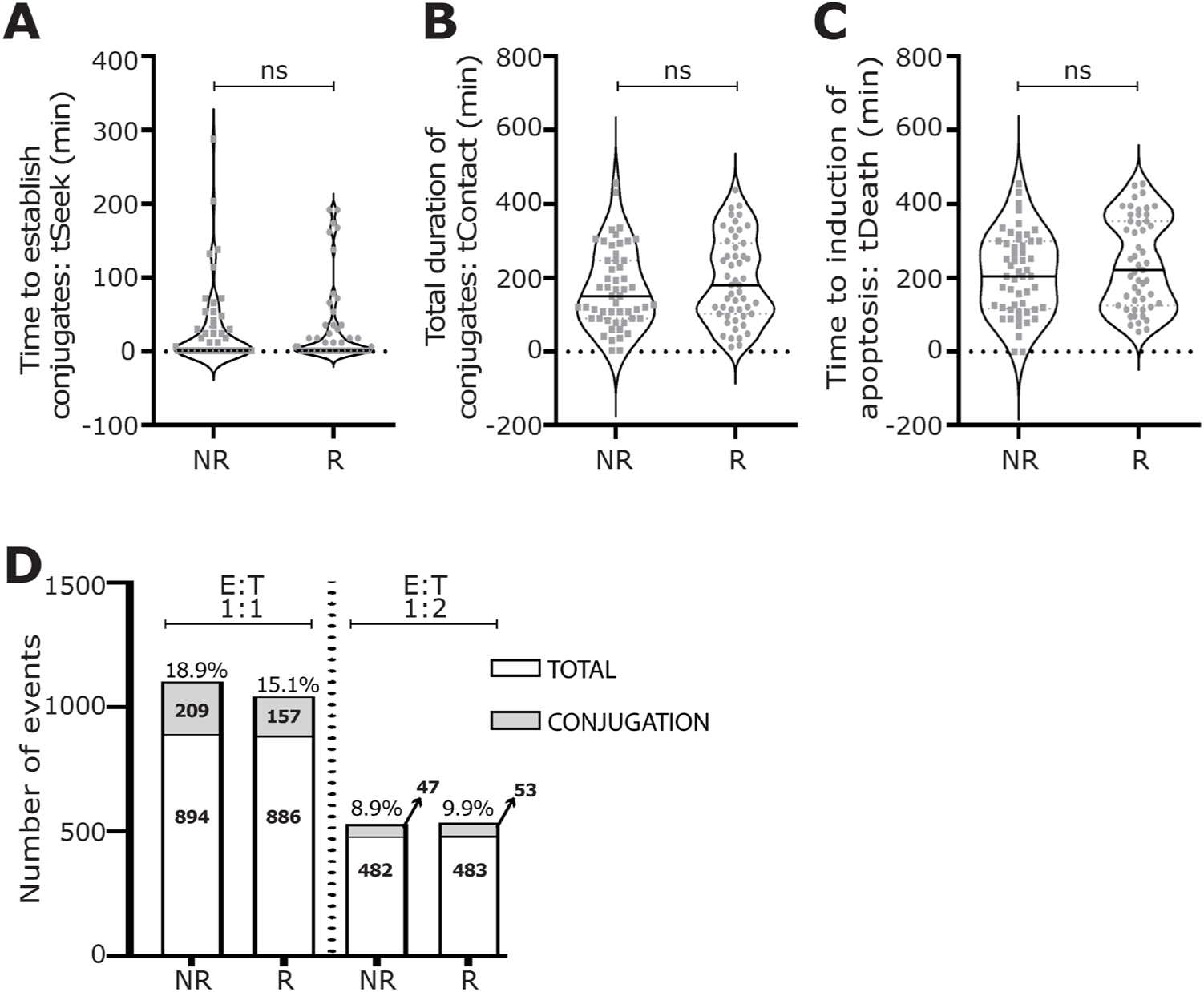
There are no significant differences in the kinetics of tumor cell killing between R and NR TILs as measured by TIMING. Stained R/NR TILs and stained autologous tumor cells were loaded on a nanowell array to profile the interactions at a single-cell level. **(A)** Comparison of the time the effector cell (TILs) established a synapse (tSeek) with the target tumor cells. **(B)** Comparative assessment of the time of cumulative conjugation (tContact) between TIL and tumor cells. **(C)** Comparison of the time to induce tumor cell apoptosis (tDeath) [annexin V (AF647)^+^ cells] between TIL and tumor cells. **(D)** Number of events in wells where we had one TIL and multiple target cells (1:1, 1:2, and 1:3) with/without conjugation between R and NR TILs. These data are from a total of 3,560 individual TILs. All data are from a set of TILs, two R and two NR, interrogated against autologous tumor cells. All data in panels A-C are from an E:T ratio of 1:1. *p* values for comparisons were computed using a non-parametric Mann-Whitney test. ns = no significance (A-C).

**Supplementary figure 7.**
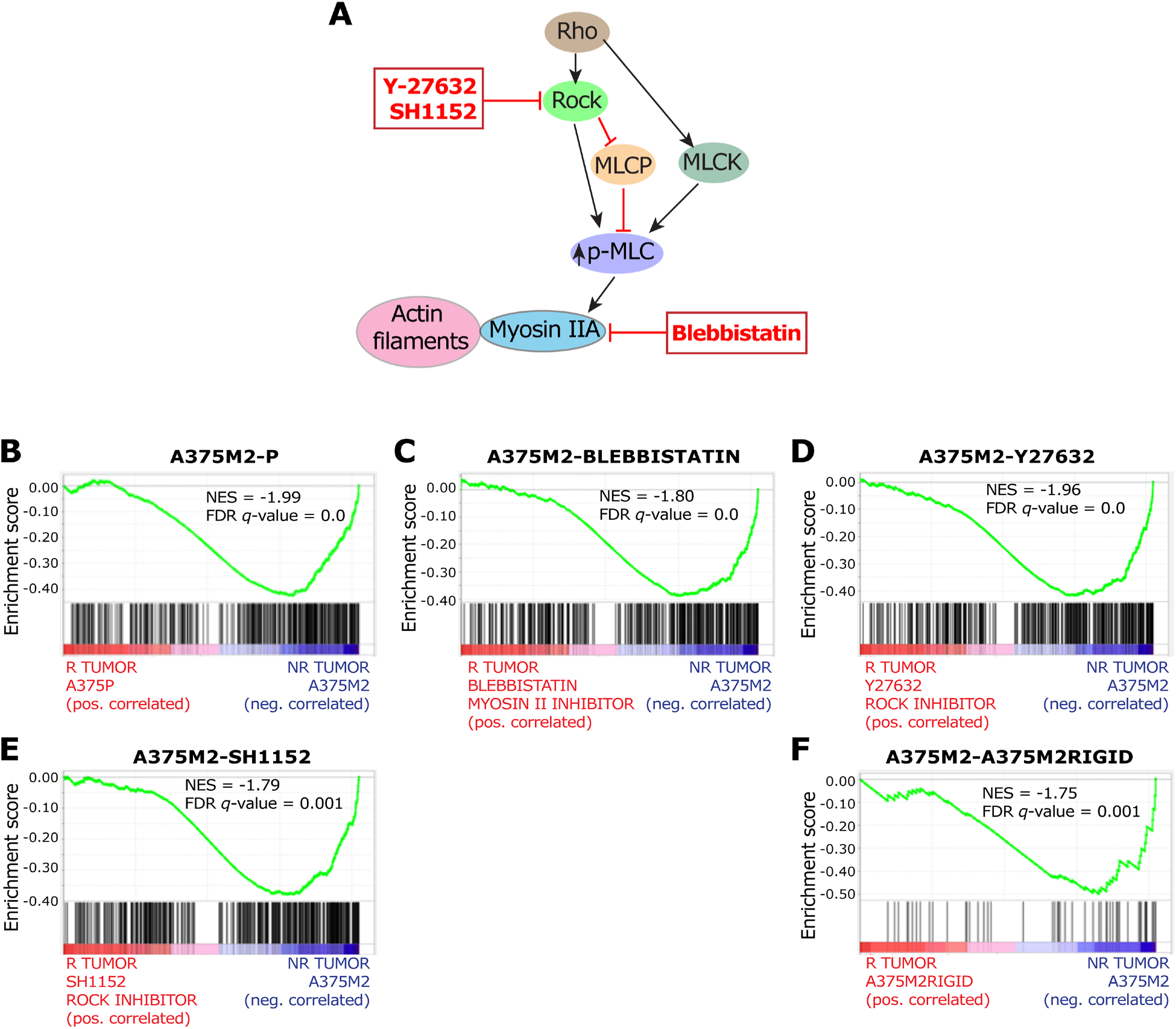
NR tumor cells were enriched in transcripts associated with amoeboid signature in comparison to R tumor cells. Signatures related to amoeboid-like motility cells treated or not with different actomyosin contractility inhibitors reported previously (Georgouli *et al*., 2019) were used to compare the R and NR tumor cells data. **(A)** The Rho-Rock-Myosin signaling pathway controls amoeboid migration and the role of pharmacological inhibitors. Rho-associated kinase (Rock), myosin light chain (MLC), MLC kinase (MLCK), and myosin light chain phosphatase (MLCP). **(B)** GSEA plots comparing R and NR tumor cells to the A375M2 (highly metastatic melanoma cell line, with higher Myosin II activity) and A375M2P (poorly metastatic cell line, with lower Myosin II activity) signatures. **(C)** GSEA plots comparing R and NR tumor cells to the A375M2 and the A375M2 treated cells with blebbistatin (inhibitor of Myosin II ATPase) signatures. (**D-E)** GSEA plots comparing R and NR tumor cells to the A375M2 and the A375M2M2 cells treated with Y27632 (D) or SH1152 (E) (contractility inhibitors of Rho-kinase protein) signatures. **(F)** GSEA plots comparing R and NR tumor cells to the A375M2 and the A375M2 cells plated on a rigid collagen matrix (interfering with the contractility of the cells) signatures. All comparative signatures are derived from GSE: GSE23764

**Supplementary figure 8.**
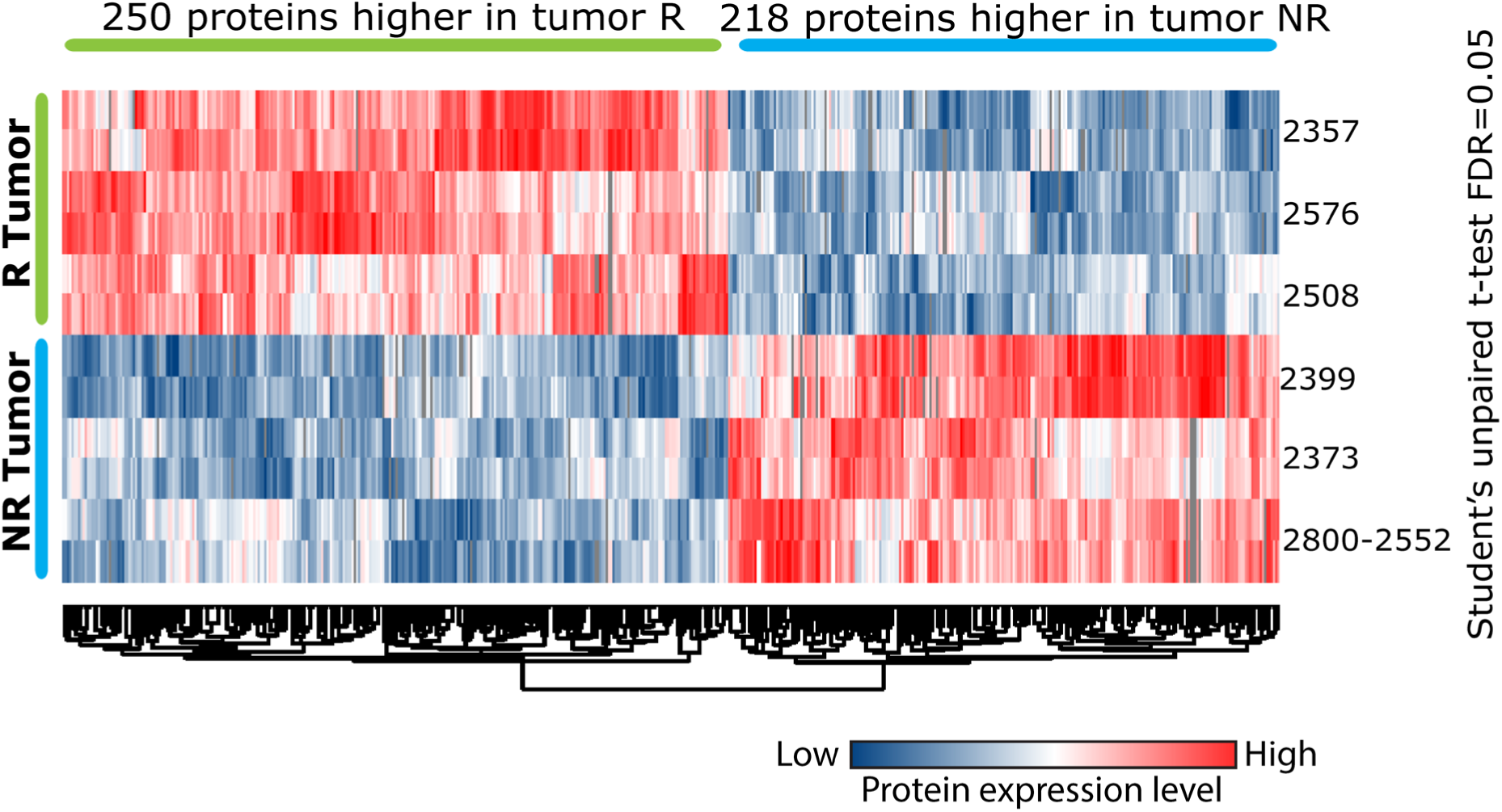
Differentially expressed proteins (DEPs) between R and NR tumor cells. Cell lysates from three R and three NR tumor cells were trypsin-digested and fractionated, followed by a high-resolution LC-MS/MS analysis. Hierarchical clustering of 468 significantly changing proteins between the two response groups (FDR < 0.05). The color indicates the Z-scored expression level of each DEP (red and blue for high and low levels, respectively).

**Supplementary figure 9.**
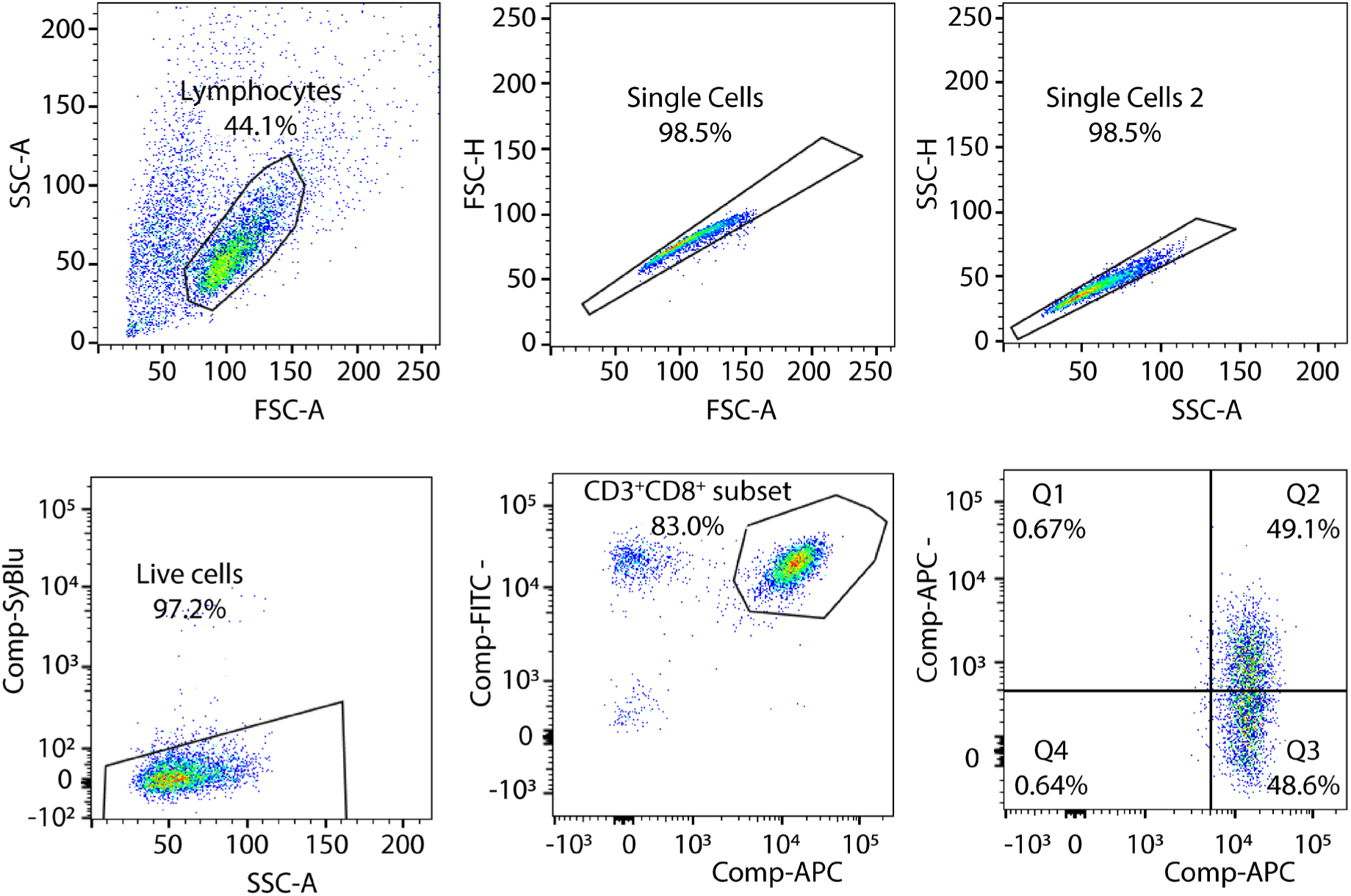
Gating strategy used to flow sort CD8^+^CD69^+^ and CD8^+^CD69^−^TILs. TILs were co-cultured with autologous tumor cells for 12 h. The cells were harvested, stained, and sorted using FACS Aria Fusion Cell Sorter (BD Bioscience), and data analysis was conducted using FlowJo software (Tree Star, Inc.). The plots indicate the gating strategy. We sorted CD3^+^CD8^+^CD69^+^ and CD3^+^CD8^+^CD69^-^ TILs from three Rs and three NRs (n = 6). Live cells were gated as pacific blue negative cells.

**Supplementary figure 10.**
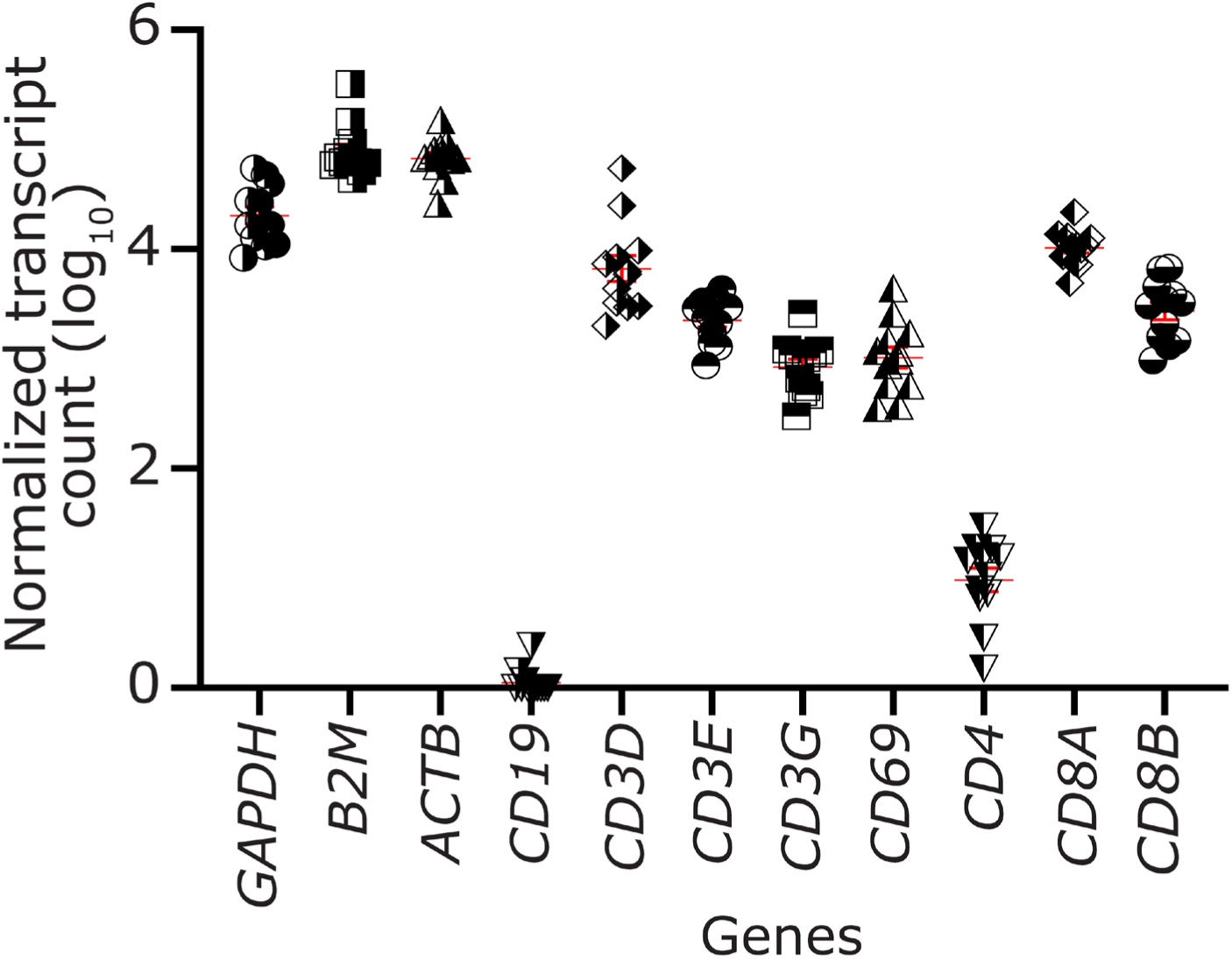
RNA-seq validation of the flow-sorted TILs. TILs were co-cultured with their autologous tumor cells for 12 hours; subsequently, cells were sorted as CD3^+^CD8^+^CD69^+^ and CD3^+^CD8^+^CD69^-^ TIL by FACS (6 R TIL and 6 NR TIL populations). cDNA libraries were prepared for RNA-seq, and the normalized transcripts count of housekeeping genes and transcripts related to CD8^+^ T-cells were analyzed as quality control of TIL RNA-seq.

**Supplementary figure 11.**
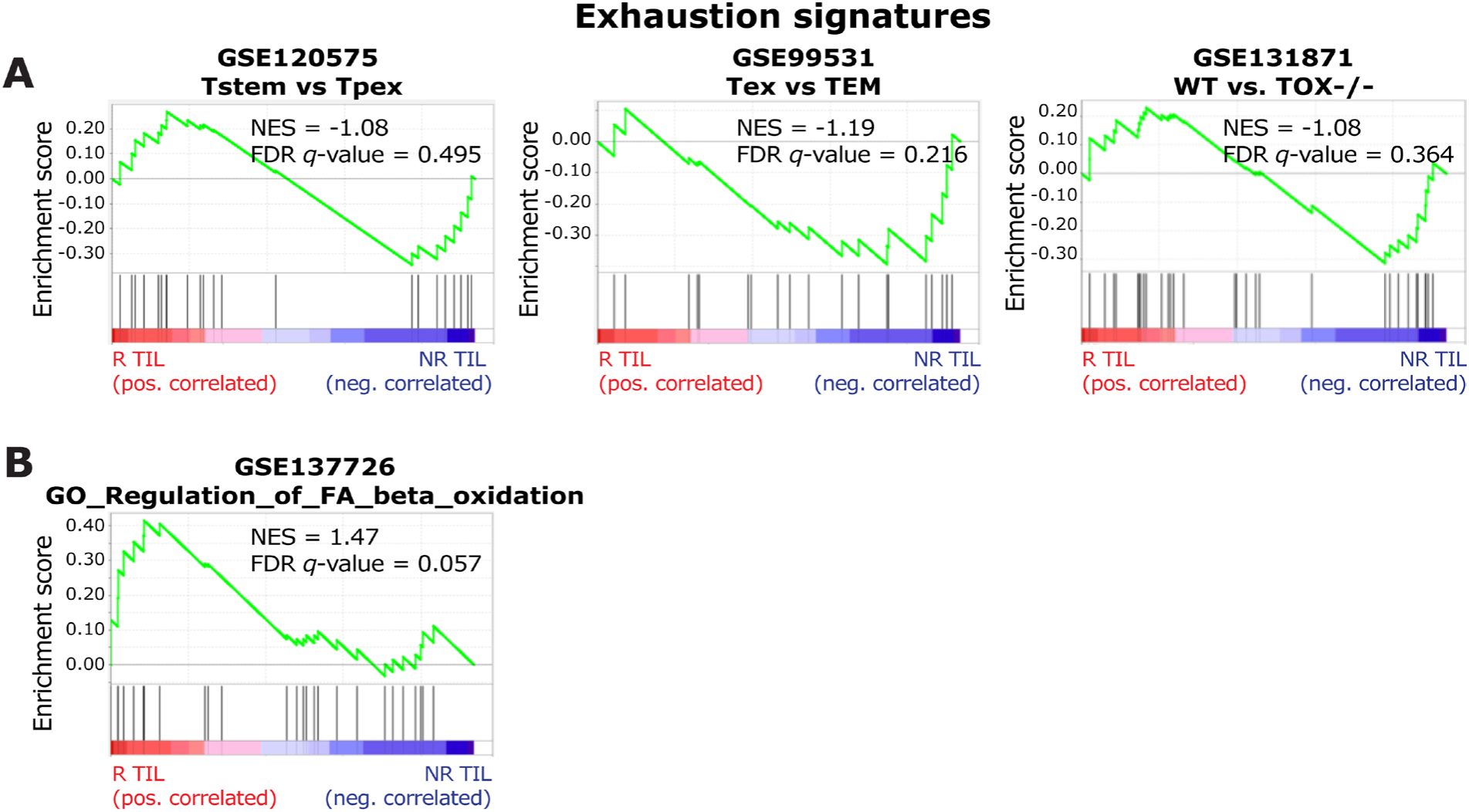
R TILs are not exhausted T-cells. **(A)** GSEA plots comparing R and NR TILs to three different published signatures of CD8^+^ T exhausted (T_ex_) and T progenitor exhausted-like (T_PEX_) [GSE120575, GSE99531, GSE131871]. **(B)** GSEA plots comparing R and NR TILs to a known signature of the FA beta-oxidation pathway (GSE137726).

**Supplementary figure 12.**
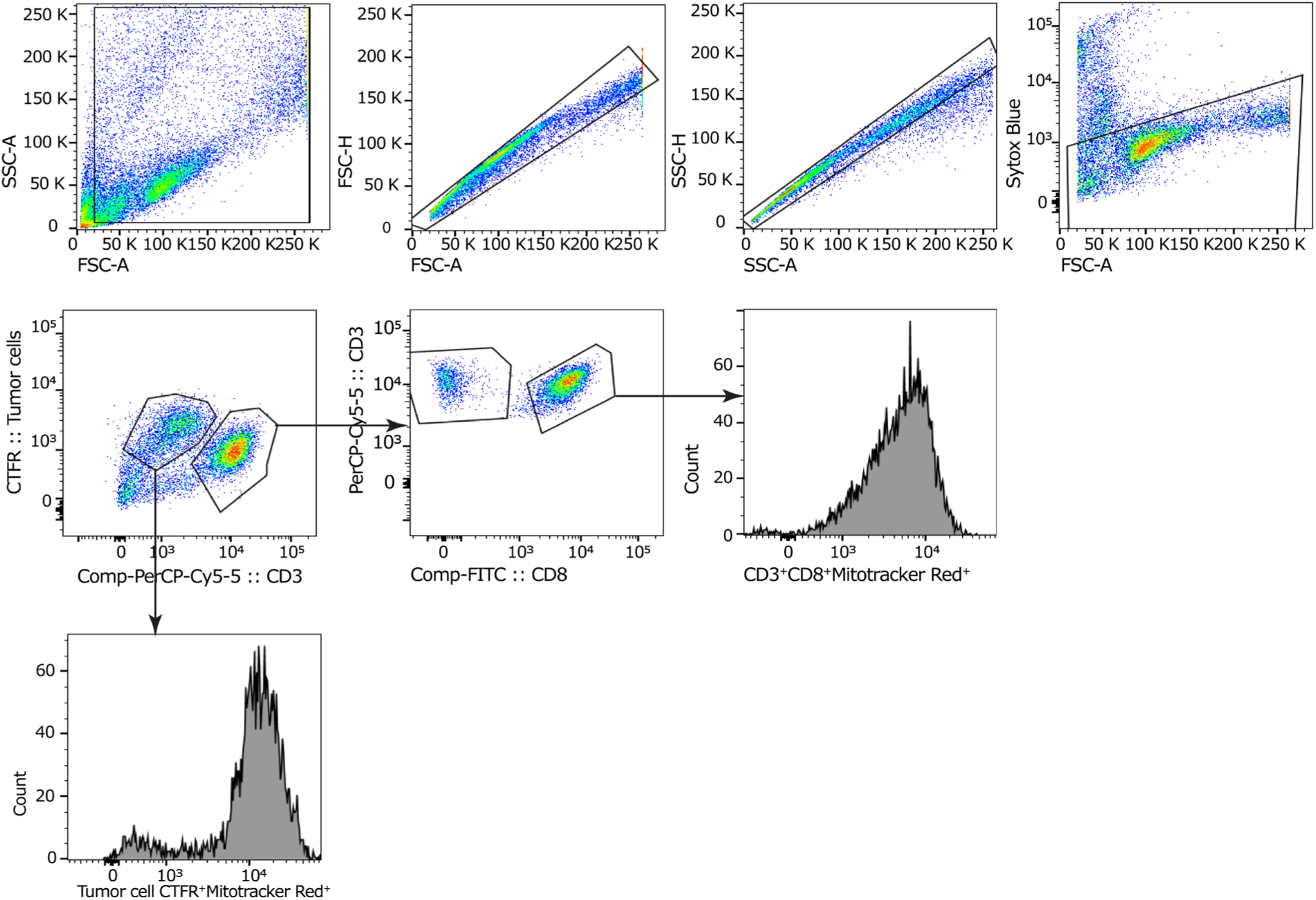
Gating strategy used to track the mitochondria, LD and FA on TILs. TILs were co-cultured with their autologous tumor cells (pre-stained with CellTrace Far Red [CTFR]) for 12 h. The cells were washed, stained with the indicated antibodies or dyes, and analyzed using a flow cytometer. TILs were gated as follows: CD3^+^CD8^+^MitoTracker green (MTG)^+^CTFR^-^, CD3^+^CD8^+^2NBDG^+^CTFR^-^, CD3^+^CD8^+^MitoTracker red (MTR)^+^CTFR^-^, CD3^+^CD8^+^Bodipy 498/503^+^ CTFR^-^, and CD3^+^CD8^+^Bodipy 558/568 (C_12_)^+^ CTFR^-^ . Dead cells were excluded based on Ghost live/dead violet 450 stainings. Data were analyzed using the LSR. Fortessa X-20 analyzer (BD Biosciences) and data analysis were conducted using FlowJo software (Tree Star, Inc.)

**Supplementary figure 13.**
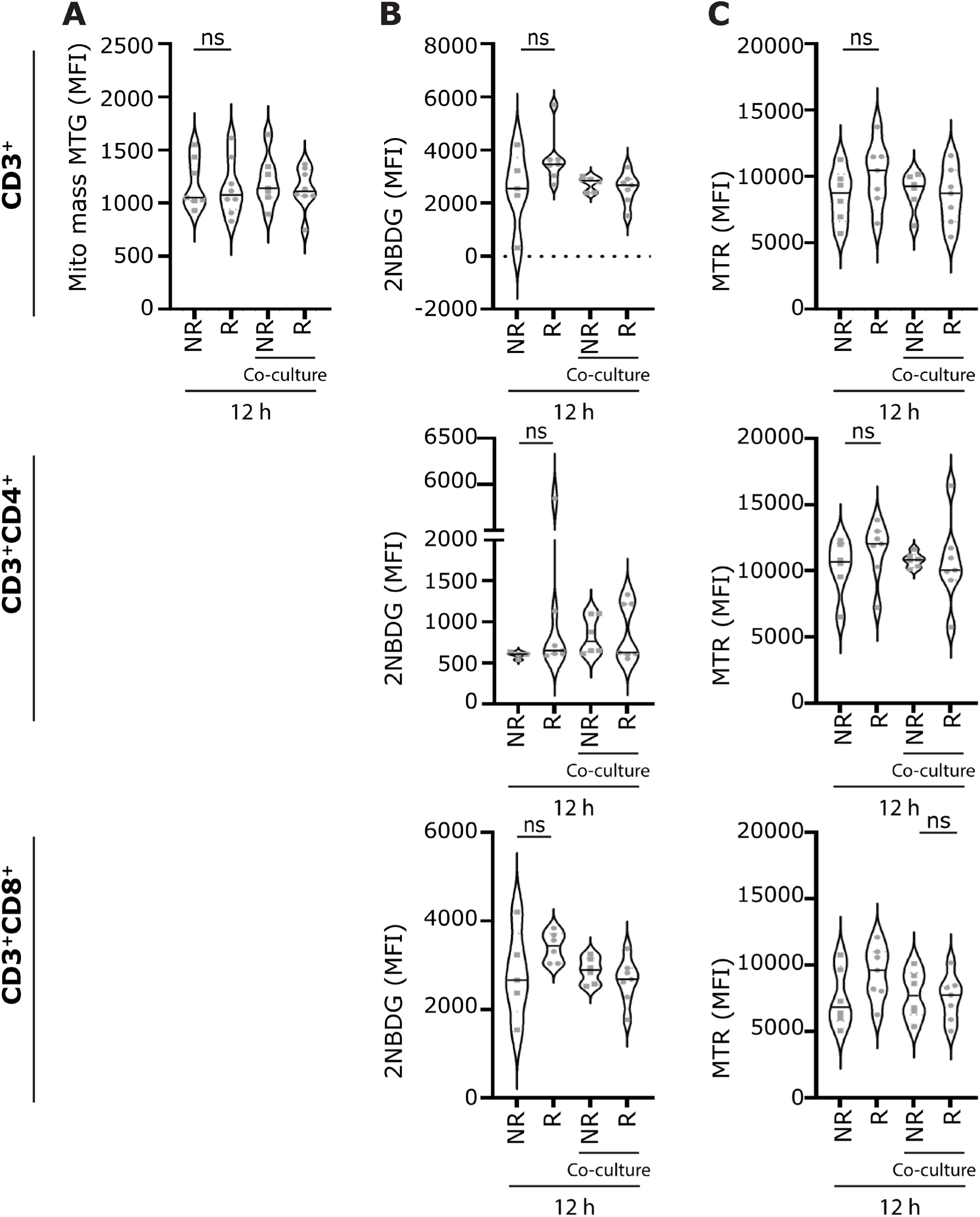
TILs show no significant alteration in glucose and mitochondria metabolism in TILs from Rs and NRs. TILs with or without co-culture with autologous tumor cells were stained with the indicated antibodies and dyes and measured by flow cytometry. The MFI analysis of the indicated dyes measured on the CD3^+^, CD3^+^CD4^+^, and CD3^+^CD8^+^ gated TILs are shown. (n = 12 [six R, ad six NR TILs]). Dead cells were excluded based on Ghost live/dead violet 450 staining. **(A)** Comparison of the mitochondria mass measured by staining TIL with MitoTracker green (MTG), **(B)** Comparative assessment of the difference in glucose uptake measured via the fluorescent glucose analog 2NBDG. **(C)** Comparison of the membrane potential measured by staining TIL with MitoTracker red CMXRos (MTR). *P* values for comparisons were computed using a nonparametric Mann-Whitney test. ns = no significant.

**Supplementary figure 14.**
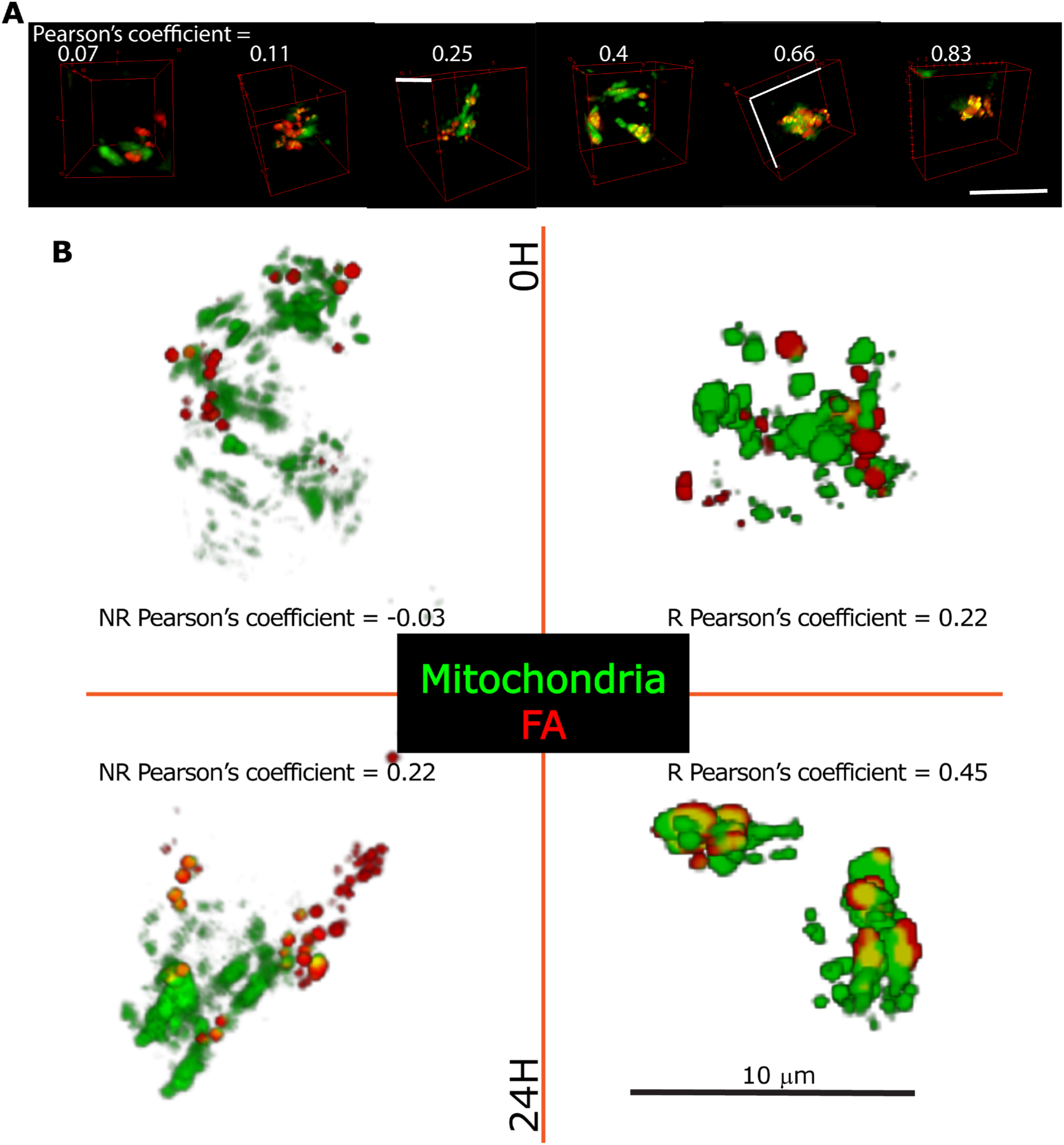
Representative examples of TILs for monitoring the uptake of fatty acids (FA) using Bodipy 558/568 (C_12_). TILs pulsed with the fluorescent FA analog Bodipy 558/568 (C_12_) for 12 hours to enable uptake of this FA (pulse). The TILs were co-cultured with their autologous tumor cells in HBSS (starvation, chase) for 24 hours. The total FA and their colocalization with mitochondria were quantified by confocal microscopy. **(A)** Representative images showing examples of different values of Pearson’s coefficient for colocalization of FA and mitochondria. Scale bar = 10 μm. **(B)** Representative composite micrographs of the R and NR TILs median Pearson’s coefficient found at 0 and 24 hours, FA stained with Bodipy 558/568 (C_12_) and mitochondria with MitoTracker green (MTG). Scale bar = 10 μm..

**Supplementary figure 15.**
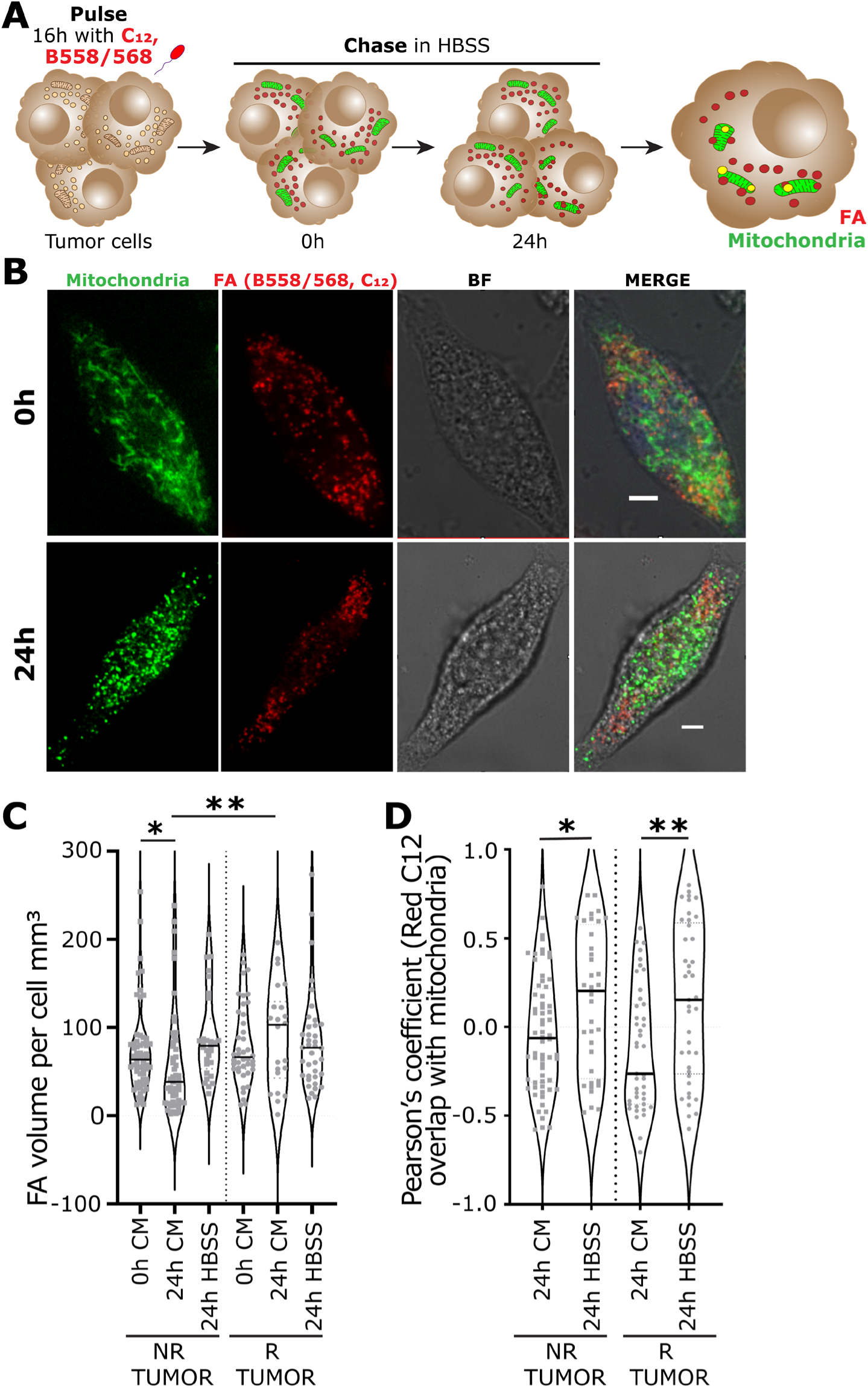
R and NR TILs did not show significant differences in the uptake of the utilization of FAs. **(A)** Tumor cells pulsed with Bodipy 558/568 (C_12_) for 16 h to enable uptake of this FA (pulse). The cells were washed, stained with MTG (mitochondria mass), and cultured in HBSS (starvation, chase). The total FA and their colocalization with mitochondria were quantified by confocal microscopy. N = 7, three R and 4 NR tumor cells. **(B)** Representative confocal images of fatty acid and mitochondria on an NR tumor cell at 0h and 24 h. Scale bar = 25 μm. BF = Brightfield. **(C)** Comparison of the volume of FA per cell between R and NR tumor cells after 24 hours in CM and starvation conditions (HBSS). These data are from a total of 243 individual tumor cells. **(D)** Comparison of the colocalization of the FA and mitochondria using Pearson’s coefficient between R and NR tumor cells after 24 hours in starvation conditions (HBSS). These data are from a total of 184 individual tumor cells. *p* values for comparisons were computed using a nonparametric Mann-Whitney test. \**p* < 0.05, \*\**p* < 0.01,\*\*\*\**p* < 0.0001 in C-D.

